# Epstein-Barr Virus Latent Membrane Protein 1 Subverts IMPDH pathways to drive B-cell oncometabolism

**DOI:** 10.1101/2024.11.07.622457

**Authors:** Eric M. Burton, Jin Hua Liang, Bidisha Mitra, John M. Asara, Benjamin E. Gewurz

**Affiliations:** Division of Infectious Diseases, Department of Medicine, Brigham and Women’s Hospital, 181 Longwood Avenue, Boston, MA 02115, USA; Center for Integrated Solutions for Infectious Diseases, Broad Institute of Harvard and MIT, Cambridge, MA 02142, USA; Department of Microbiology, Harvard Medical School, Boston, MA 02115, USA; Harvard Program in Virology, Boston, MA 02115, USA; Division of Signal Transduction, Beth Israel Deaconess Medical Center and Department of Medicine, Harvard Medical School, Boston, MA, United States; Harvard Program in Virology, Harvard Medical School, Boston, MA 02115, USA; Harvard Program in Virology, Boston, MA 02115, USA

## Abstract

Epstein-Barr virus (EBV) is associated with multiple types of cancers, many of which express the key viral oncoprotein Latent Membrane Protein 1 (LMP1). LMP1 is the only EBV-encoded protein whose expression is sufficient to transform both epithelial and B-cells. Although metabolism reprogramming is a cancer hallmark, much remains to be learned about how LMP1 alters lymphocyte oncometabolism. To gain insights into key B-cell metabolic pathways subverted by LMP1, we performed systematic metabolomic analyses on B cells with conditional LMP1 expression. This approach highlighted that LMP highly induces *de novo* purine biosynthesis, with xanthosine-5-P (XMP) as one of the most highly LMP1-upregulated metabolites. Consequently, IMPDH inhibition by mycophenolic acid (MPA) triggered apoptosis of LMP1-expressing EBV-transformed lymphoblastoid cell lines (LCL), a key model for EBV-driven immunoblastic lymphomas. Whereas MPA instead caused growth arrest of Burkitt lymphoma cells with the EBV latency I program, conditional LMP1 expression triggered their apoptosis. Although both IMPDH isozymes are expressed in LCLs, only IMPDH2 was critical for LCL survival, whereas both contributed to proliferation of Burkitt cells with the EBV latency I program. Both LMP1 C-terminal cytoplasmic tail domains critical for primary human B-cell transformation were important for XMP production, and each contributed to LMP1-driven Burkitt cell sensitivity to MPA. MPA also de-repressed EBV lytic antigens including LMP1 in latency I Burkitt cells, highlighting crosstalk between the purine biosynthesis pathway and the EBV epigenome. These results suggest novel oncometabolism-based therapeutic approaches to LMP1-driven lymphomas.

**IMPORTANCE:** Altered metabolism is a hallmark of cancer, yet much remains to be learned about how EBV rewires host cell metabolism to support multiple malignancies. While the oncogene LMP1 is the only EBV-encoded gene that is sufficient to transform murine B-cells and rodent fibroblasts, knowledge has remained incomplete about how LMP1 alters host cell oncometabolism to aberrantly drive infected B-cell growth and survival. Likewise, it has remained unknown whether LMP1 expression creates metabolic vulnerabilities that can be targeted by small molecule approaches to trigger EBV-transformed B-cell programmed cell death. We therefore used metabolomic profiling to define how LMP1 signaling remodels the B-cell metabolome. We found that LMP1 upregulated purine nucleotide biosynthesis, likely to meet increased demand. Consequently, LMP1 expression sensitized Burkitt B-cells to growth arrest upon inosine monophosphate dehydrogenase blockade. Thus, while LMP1 itself may not be a therapeutic target, its signaling induces dependence on downstream druggable host cell nucleotide metabolism enzymes, suggesting rational therapeutic approaches.

## Introduction

Epstein-Barr virus (EBV) persistently infects over 90% of adults worldwide. EBV causes infectious mononucleosis, is a key multiple sclerosis trigger and contributes to 200,000 cancers per year. EBV is associated with a wide range of lymphomas, including endemic Burkitt lymphoma (BL), Hodgkin lymphoma, post-transplant lymphoproliferative diseases (PTLD), T and NK cell lymphomas. EBV is also highly associated with nasopharyngeal carcinoma and gastric carcinoma^1–6^.

To establish lifelong colonization of the memory B cell compartment, EBV uses a series of latency programs, in which combinations of oncogenic Epstein-Barr nuclear antigens (EBNA) and latent membrane proteins (LMP) are expressed. The highly B-cell transforming latency III program is comprised LMP1 and LMP2A, all six EBNA and non-coding RNAs^7–9^. LMP1 and LMP2A mimic signaling from activated CD40 and B-cell receptors, respectively. Latency III is hypothesized to drive infected B-cells into secondary lymphoid germinal centers, within which infected B-cells switch to the latency II program, comprised of EBNA1, LMP1 and LMP2A. Germinal center cytokines boost LMP1 expression through JAK/STAT signaling^10,11^. Upon memory B-cell differentiation, EBV switches again to the latency I program, in which EBNA1 is the only EBV-encoded protein expressed. Most Burkitt lymphomas express latency I^6,12–14^.

LMP1 is the only EBV-encoded oncogene whose expression is sufficient to transform B lymphocytes and epithelial cells^15–21^. LMP1 is comprised of a 24 residue N-terminal cytoplasmic tail, six transmembrane (TM) domains and a 200 residue C-terminal cytoplasmic tail ^8,9,22,23^. LMP1 TM domains drive lipid raft association and constitutive signaling by cytoplasmic tail regions ^24–26^. Two LMP1 C-terminal tail regions, termed C-terminal activating region (CTAR) or transformation effector site (TES), are essential for primary human B-cell transformation. CTAR1/TES1 activates canonical and non-canonical NF-κB, PI3 kinase, MAP kinase (MAPK) and JAK/STAT signaling^8,9,27–31^, whereas CTAR2/TES2 activates canonical NF-κB, MAP, IRF7 and P62 pathways ^8,9,23,24,32–39^. The LMP1 CTAR3 region activates JAK/STAT and SUMOylation pathways ^40–42^.

EBV oncogenes have not yet proven to be druggable *in vivo*, though induce downstream host cell dependencies that may instead be attractive therapeutic targets, including within remodeled oncometabolism pathways. EBV-driven metabolic remodeling is necessary for primary human B cell transformation into immortalized lymphoblastoid cell lines (LCL), a major PTLD model^43–46^. For instance, EBV manipulates nucleotide metabolism to support oncogene-driven demand^47–49^, in which infected cells become highly dependent upon EBV upregulated nucleotide metabolism pathways, including one-carbon metabolism^45^. EBNA2 and LMP1 jointly induce the *de novo* cytidine biosynthesis pathway, which then exerts critical roles in EBV-transformed B-cell proliferation^43^. However, while metabolism reprogramming is a hallmark of cancer^50^, LMP1 effects on host B-cell metabolism pathways remain to be fully characterized.

Here, we use metabolomic approaches to characterize LMP1 roles in host B-cell oncometabolism remodeling. Liquid chromatography/mass spectrometry (LC/MS) profiling identified that LMP1 reprogrammed a wide range of metabolic pathways, in particular purine metabolism. LMP1 upregulated levels of the guanosine monophosphate precursor xanthosine-5-phosphate, which is produced by activity of the enzymes inosine monophosphate dehydrogenase 1 and 2 (IMPDH1/2). IMPDH blockade by mycophenolic acid (MPA) or by CRISPR editing triggered LCL apoptosis, which was rescuable by guanosine triphosphate GTP supplementation. LMP1 sensitized latency I Burkitt cells to apoptosis upon MPA treatment, and this phenotype was dependent upon TES1 and TES2 signaling. Within latency I, IMPDH blockade triggered growth arrest and reshaped the latency I EBV epigenome.

## Results

### Metabolomic analyses highlights key pathways targeted by LMP1

To gain insights into LMP1 B-cell oncometabolism remodeling roles, we constructed Burkitt lymphoma B-cell lines with conditional LMP1 expression. We validated that doxycycline-induced LMP1 expression triggered NF-κB pathway activation and LMP1 target gene expression in EBV+ Daudi EBV+ as well as in EBV-negative Akata Burkitt cells (**Figure S1A-D**). We then performed polar metabolite LC/MS profiling of both Daudi and Akata cell models, mock induced or induced for LMP1 expression for 24 hours by doxycycline addition (**Figure 1A**). LMP1 significantly induced 124 and reduced 10 Daudi metabolites (**Figure 1B**, **Table 1**). Similarly, LMP1 induced 56 and reduced 11 Akata metabolites (**Figure 1C**, **Table 1**). Metabolite pathway impact analysis^51^ identified that purine metabolism was amongst the most highly LMP1-induced metabolic pathway in both Burkitt model systems (**Figure 1D, E**).

**Figure 1.**
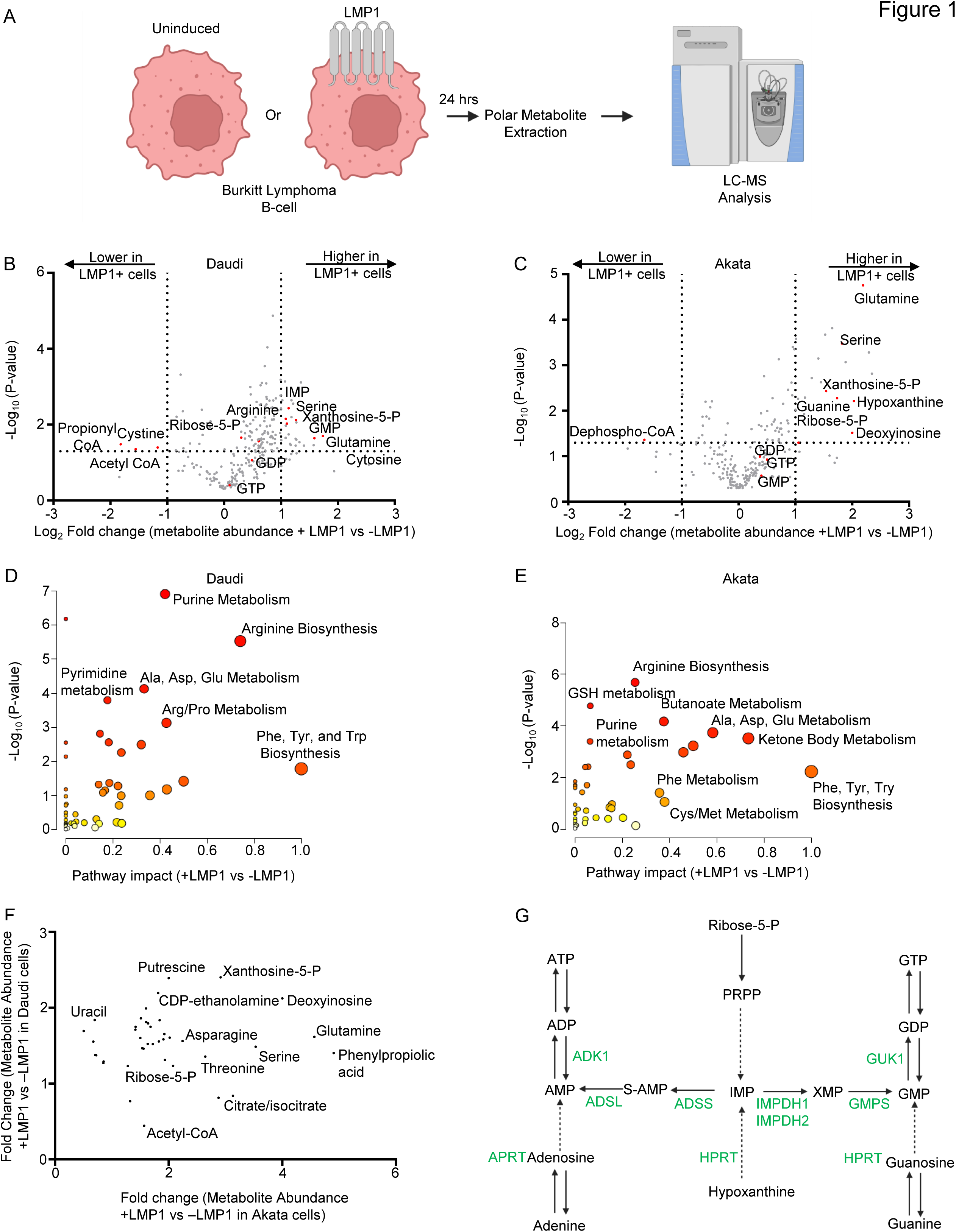
LMP1-mediated B-cell metabolome remodeling. (A) Metabolomic experiment workflow. Conditional LMP1 Daudi or Akata Burkitt cells were mock induced or induced by doxycycline (250ng/ml) for LMP1 expression for 24h. Polar metabolites were analyzed by targeted metabolomic analysis. (B) Volcano plot analysis of liquid chromatography mass spectrometry (LC-MS) analysis of n=3 replicates of Daudi cells mock induced or induced for LMP1 expression. Positive fold changes indicate higher metabolite concentrations in LMP1+ vs LMP1-cells. Selected host cell metabolites induced vs. suppressed by LMP1 are indicated. (C) Volcano plot analysis of LC-MS analysis of n=6 replicates of Akata cells mock induced or induced for LMP1 expression. (D) Metaboanalyst analysis of LMP1 driven Daudi cell metabolic pathway impact from the data presented in (B), using data from significantly changed metabolites in LMP1+ vs LMP1-cells (*p* value > 0.05). Higher pathway impact values indicates stronger effects of conditional LMP1 expression on the indicated pathway. Purine metabolism was amongst the most highly LMP1 impacted pathways. (E) Metaboanalyst pathway impact analysis of LMP1+ vs LMP1-Akata cells from the data presented in (C). (F) Volcano plot analysis cross-comparing fold change of metabolite abundances in LMP1+ versus LMP1-Akata cells (x-axis) vs Daudi cells (Y-axis). Shown are metabolites whose abundances were LMP1 increased by ≥1.2 fold in both cell models. Selected metabolites are annotated, including xanthosine-5-phosphate, which was highly LMP1-induced under both conditions. (G) Purine metabolism pathways. The *de novo* pathway uses Ripose-5-phosphate and PRPP to generate inosine monophosphate (IMP), whereas the salvage pathway metabolizes hypoxanthine into IMP. IMP can be converted by IMPDH1/2 to xanthosine monophosphate (XMP) and subsequently to guanosine monophosphate (GMP). Alternatively, IMP can be donverted to adenosine monophosphate (AMP).

The purine biosynthesis metabolite xanthsoine-5-phosphate (XMP), produced from the precursor inosine monophosphate by the enzymes inosine monophosphate dehydrogenase (IMPDH) 1 and 2, was the most highly LMP1 upregulated metabolite in both Akata and Daudi cells (**Figure 1F-G**, **Table 2**). XMP is converted into guanosine monophosphate (GMP), guanosine diphosphate (GDP) and guanosine triphosphate (GTP) (**Figure 1G)**. However, LMP1 more modestly increased GMP, GDP and GTP levels, suggesting that LMP1 may also increase their consumption. Further underscoring LMP1 nucleotide metabolism remodeling, LMP1 increased steady state levels of the key purine nucleotide biosynthetic building block ribose-5-phosphate (**Figure 1F-G**). LMP1 also increased levels of the hypoxanthine precursor deoxyinosine, suggesting that LMP1 may also support purine demand through upregulation of purine salvage metabolism (**Figure 1F-G)**.

LMP1 also highly induced amino acid metabolism in both Daudi and Akata. Pathway impact analysis^51^ highlighted that glutamine metabolism was highly induced in both Burkitt models (**Figure 1D, E**). Notably, glutamine plays major roles in *de novo* purine and pyrimidine synthesis and was the most highly LMP1 induced amino acid in Akata and Daudi (**Fig. 1F**). LMP1 also significantly increased the abundance of serine, which is a major one carbon metabolism donor for *de novo* purine biosynthesis, including in newly EBV-infected and EBV-transformed LCLs^45^. Together, these metabolomic profiling results suggest that LMP1 remodels B-cell nucleotide and amino acid metabolism pathways to support nucleotide demand.

### Latency III induces IMPDH metabolism dependency

Given that XMP was the most highly LMP1-induced metabolite, we next investigated the effects of inhibition IMPDH1/2 inhibition by the highly selective antagonist MPA^52,53^(**Figure 2A**). We first cross-compared MPA effects on proliferation of LMP1-negative Burkitt cells with that of two LCLs that express LMP1 as part of the latency III program. Using the carboxyfluorescein succinimidyl ester (CFSE) dye-dilution assay, in which each cell cycle results in a 50% reduction of CFSE signal, we observed that MPA inhibited Burkitt and LCL proliferation in a dose-dependent manner. This result suggests that IMPDH activity is likely necessary for maintenance of Burkitt and LCL GTP pools. By contrast, FACS analysis of vital dye 7-Aminoactinomycin D (7-AAD) uptake revealed that MPA rapidly triggered LCL but not Burkitt cell death (**Figure 2C, S2B**). We therefore cross-compared isogenic Burkitt cells that differ by EBV latency program. Consistent with our LCL results, MPA triggered cell death of latency III MUTU III cells^54^ to a significantly greater extent than latency I MUTU I (**Figures 2D, S2C**). Similarly, MPA triggered cell death to a greater extent in latency III Jijoye Burkitt cells than in its P3HR-1 subclone^55^, which harbors an EBNA2 deletion and exhibits a more restrictive form of EBV latency (**Figure 2E, S2D**). These results suggest that latency III creates a metabolic dependency on IMPDH activity for survival.

**Figure 2.**
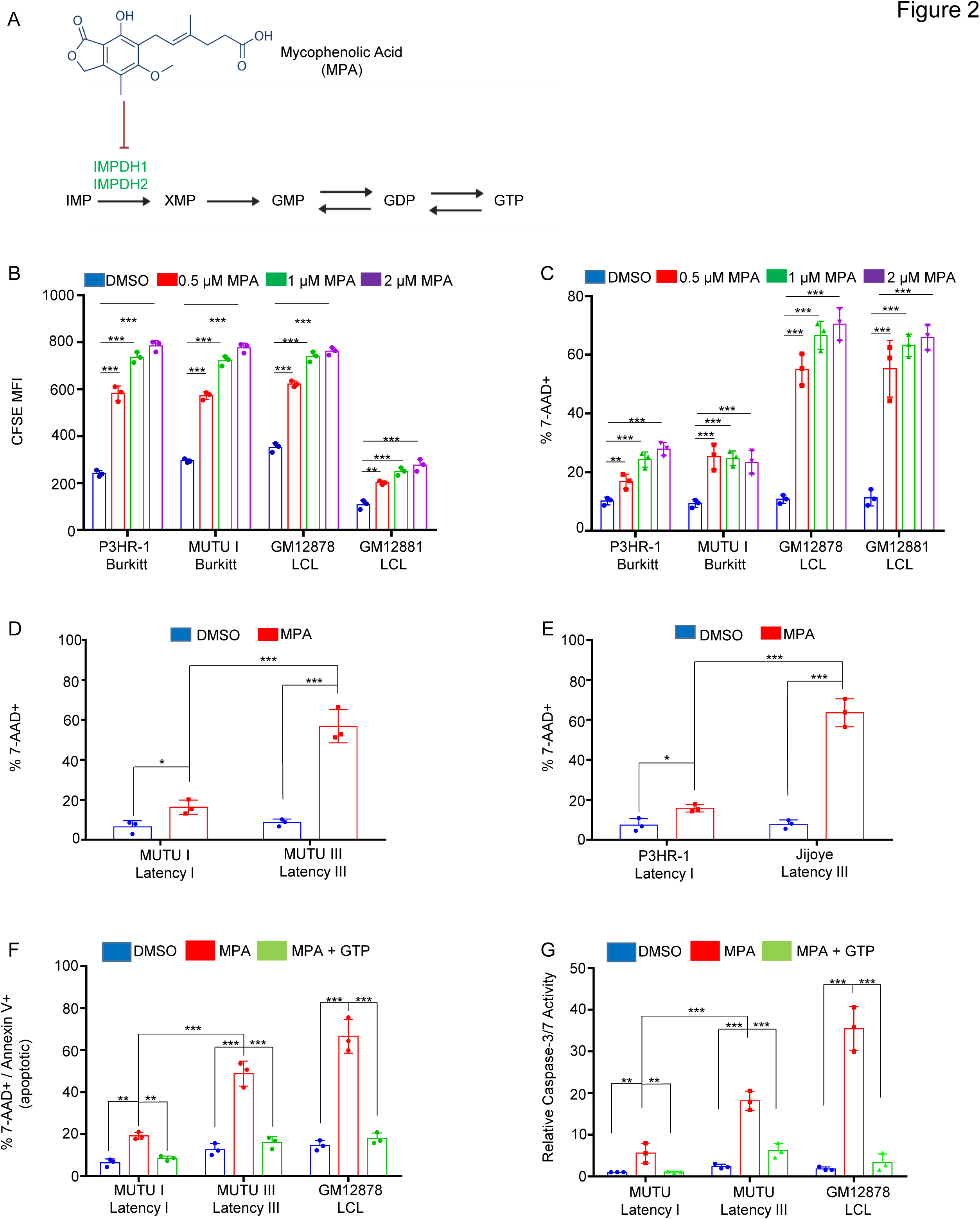
EBV latency III creates dependency on IMPDH activity for prevention of apoptosis. (A) Schematic diagram of mycophenolic acid (MPA) inhibition of the guanylate biosynthesis pathway enzymes IMPDH1 and IMPDH2. IMP, inosine monophosphate. XMP, xanthosine monophosphate, GMP, guanosine monophosphate. GDP, guanosine diphosphate. GTP, guanosine triphosphate. (B) FACS analysis of dose-dependent MPA effects on proliferation of latency I P3HR-1 and MUTU I cells versus latency III GM12878 and GM12881 LCLs, as judged by CFSE dye-dilution analysis. CFSE-stained cells were incubated with DMSO vehicle vs the indicated MPA concentrations for 96 hours and CFSE mean fluorescence intensity (MFI) was analyzed by FACS. CFSE levels are reduced by half with each mitosis. Shown are mean ± SD CFSE levels from n=3 independent replicates. (C) FACS analysis of dose-dependent effects of MPA treatment for 48 hours on cell death of latency I P3HR-1 and MUTU I cells versus latency III GM12878 and GM12881 LCLs, as judged by uptake of the vital dye 7-AAD. Shown are mean ± SD percentages of 7-AAD+ (non-viable) cells from n=3 independent replicates. (D) FACS analysis of DMSO versus MPA effects on viability of isogenic MUTU I versus III cells that differ by EBV latency I versus III programs, respectively. Shown are mean ± SD percentages of 7-AAD+ cells following DMSO versus 1 μM MPA treatment for 48 hours. (E) FACS analysis of DMSO versus MPA effects on P3HR-1 versus Jijoye Burkitt cell viability following DMSO versus 1 μM MPA treatment for 48 hours. Mean ± SD 7-AAD+ cell percentages from n=3 replicates are shown. (F) FACS analysis of mean ± SD percentages of 7-AAD+/annexin V+ cells following treatment with DMSO, 1 μM MPA with or without 100 μM GTP rescue for 48 hours. Double 7-AAD/annexin V positivity indicates apoptosis. (G) Relative caspase 3/7 activity levels of cells analyzed in panel (F), as judged by Caspase3/7 Glo assay. Mean ± SD values from n=3 replicates are shown. *, *P <* 0.05; **, *P <* 0.05; ***, *P <* 0.005; ns, nonsignificant using Student’s t-test.

Guanylate pool depletion by MPA can trigger apoptosis in cancer cells by inducing nucleotide imbalance and DNA damage^56,57^. We therefore tested whether MPA selectively increased DNA damage in latency III B-cells, using phosphorylation of the kinases ATM and ATR as readouts of host cell responses to double stranded versus single stranded DNA breakage, respectively. Unexpectedly, MPA lowered ATM and ATR phosphorylation levels (**Figure S2E**). Yet, MPA triggered apoptosis in MUTU III and GM12878 to a greater extent than in MUTU I, and this could be rescued by GTP supplementation, supporting the hypothesis that latency III drives an increased dependence on GTP pools for survival (**Figure 2F, S3**). Furthermore, MPA more highly induced executioner caspase 3 and 7 activity in MUTU III and GM12878, and this was again largely rescuable by GTP supplementation (**Figure 2G**). Taken together, our results are consistent with a model in which latency III increases both GTP demand and GTP biosynthesis, the latter of which is increased at the level of IMPDH activity.

### Both IMPDH isoforms are important for LCL proliferation

EBV induces IMPHD1 and IMPDH2 expression at the mRNA and protein levels by day two post primary human B-cell infection (**Fig. S4A-C**), a timepoint at which EBNA2 is highly expressed but before LMP1 is induced^45,58–60^. Similarly, withdrawal of conditional EBNA2 expression from 2-2-3 LCLs reduced protein levels of both IMPDH1 and IMPDH2 (**Fig. S4D**), whereas conditional Burkitt LMP1 expression did not increase IMPDH1 or 2 protein levels (**Fig. S4E**). Furthermore, IMPDH1/2 mRNA levels were not significantly depleted by 24 hours post LMP1 CRISPR knockout (KO) in GM12878 LCLs^61^, consistent with a recent report that EBNA2 and MYC but not LMP1 induce IMPDH2^45,58–60^. Therefore, whereas EBNA2 induces IMPDH1/2 expression, LMP1 may instead increase their activity.

To define IMPDH1 versus IMPDH2 roles in latency I Burkitt versus latency III LCL proliferation, we tested effects of their CRISPR KO. While depletion of IMPDH1 or IMPDH2 alone did not significantly alter MUTU I or Daudi Burkitt proliferation (**Figure 3A-B S5A-B**), depletion of either significantly impaired proliferation of the LCLs GM12878 and GM13111, with a stronger IMPDH2 KO phenotype (**Figure 3C-D, S5A-B**). Whereas IMPDH2 KO significantly reduced GM12878 LCL live cell numbers, combined IMPDH1 and 2 depletion was required to reduce P3HR-1 Burkitt viability (**Figure 3E-F, S5C**). By contrast, combined IMDPH1/2 depletion had only slightly greater effect on GM12878 viability than IMPDH2 depletion alone, as judged by 7-AAD uptake and Cell Titer Glo assays (**Figure 3E-F, S5D-E**). These data are consistent with a model in which LMP1 increases IMPDH2 activity to support elevated GTP demand in latency III.

**Figure 3.**
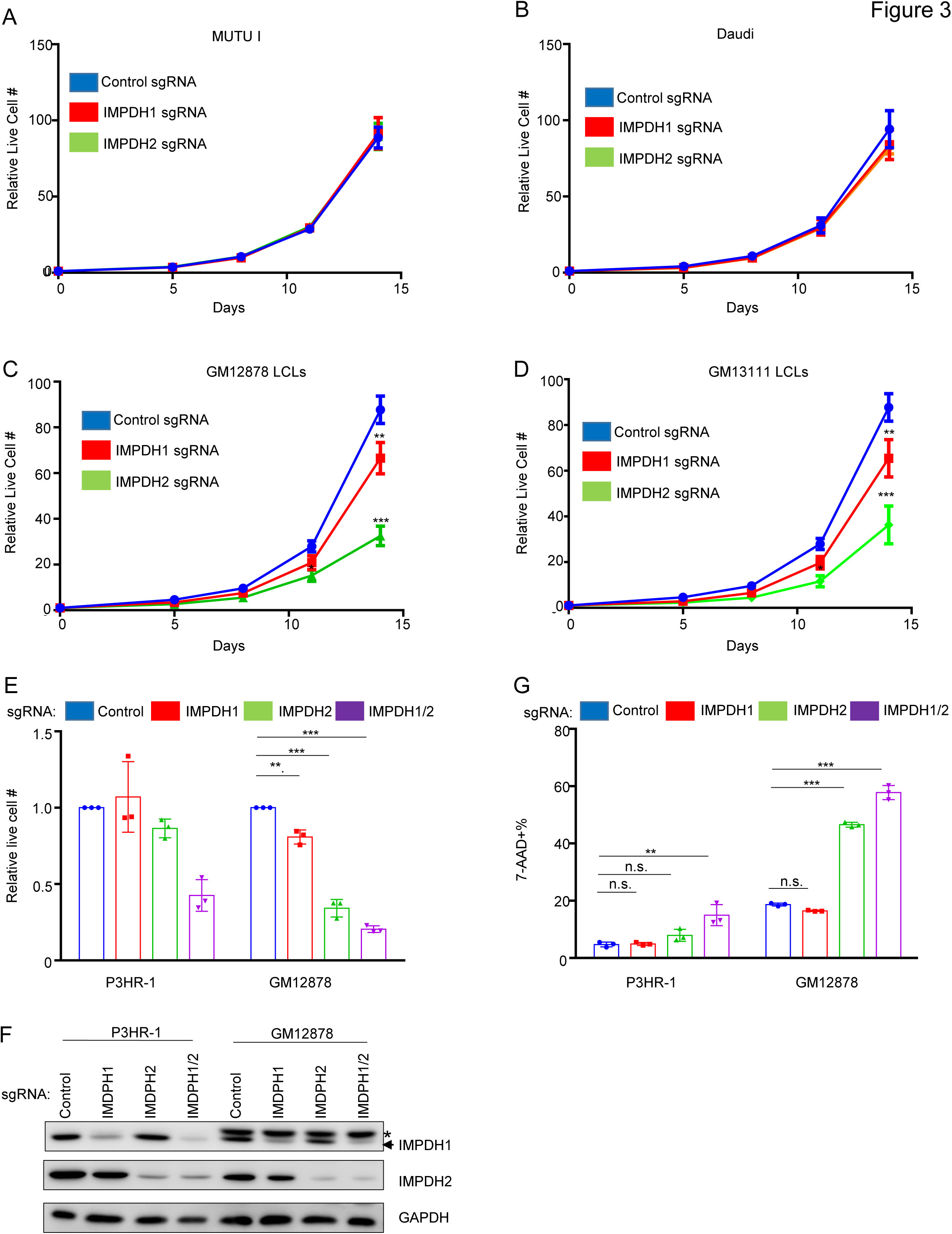
LCLs but not Burkitt cells are dependent on IMPDH2 for growth and survival. (A) Mean ± SD live cell numbers of Cas9+ MUTU I expressing control, IMPDH1 or IMPDH2 targeting single guide RNAs (sgRNA) from n=3 replicates. Cells transduced with lentiviruses expressing the indicated sgRNAs were puromycin selected. Cell numbers immediately following puromycin selection (defined as day 0 of the graph) were set to 1. Live cell numbers were quantitated by CellTiter-Glo assay. (B) Mean ± SD live cell numbers of Cas9+ Daudi cells as in (A). (C) Mean ± SD live cell numbers of Cas9+ GM12878 LCLs as in (A). (D) Mean ± SD live cell numbers of Cas9+ GM13111 LCLs as in (A). (E) Mean ± SD live cell numbers of Cas9+ P3HR-1 or GM12878 cells transduced with lentivirus expressing control, IMPDH1, IMPDH2 or IMPDH1 and 2 sgRNAs at 8 days post-puromycin selection. (F) Immunoblot analysis of WCL from Cas9+ P3HR-1 or GM12878 expresing the indicated sgRNA. *=non-specific band present in analysis of GM12878 lysates. Immunoblots are representative of n=3 independent replicates. (G) Mean ± SD MFI of Cas9+ P3HR-1 or GM12878 cells transduced with lentivirus expressing control, IMPDH1, IMPDH2 or IMPDH1 and 2 sgRNAs at 8 days post-puromycin selection, performed on cells from the same replicates shown in (E).

### LMP1 sensitizes Burkitt cells to MPA-induced apoptosis in a GTP-dependent manner

We next tested the hypothesis that LMP1 itself creates IMPDH dependency. We conditionally expressed LMP1 in Daudi cells, in the absence or presence of MPA. LMP1 signaling was not blocked by MPA, as judged by processing of the non-canonical NF-κB pathway precursor p100 into the active p52 product (**Figure 4A**). MPA induced significantly higher cell death levels in LMP1+ than in LMP1-Daudi cells, as judged by 7-AAD uptake (**Figures 4B-C**), indicating that LMP1 signaling creates a metabolic dependence on IMPDH activity for B-cell survival.

**Figure 4.**
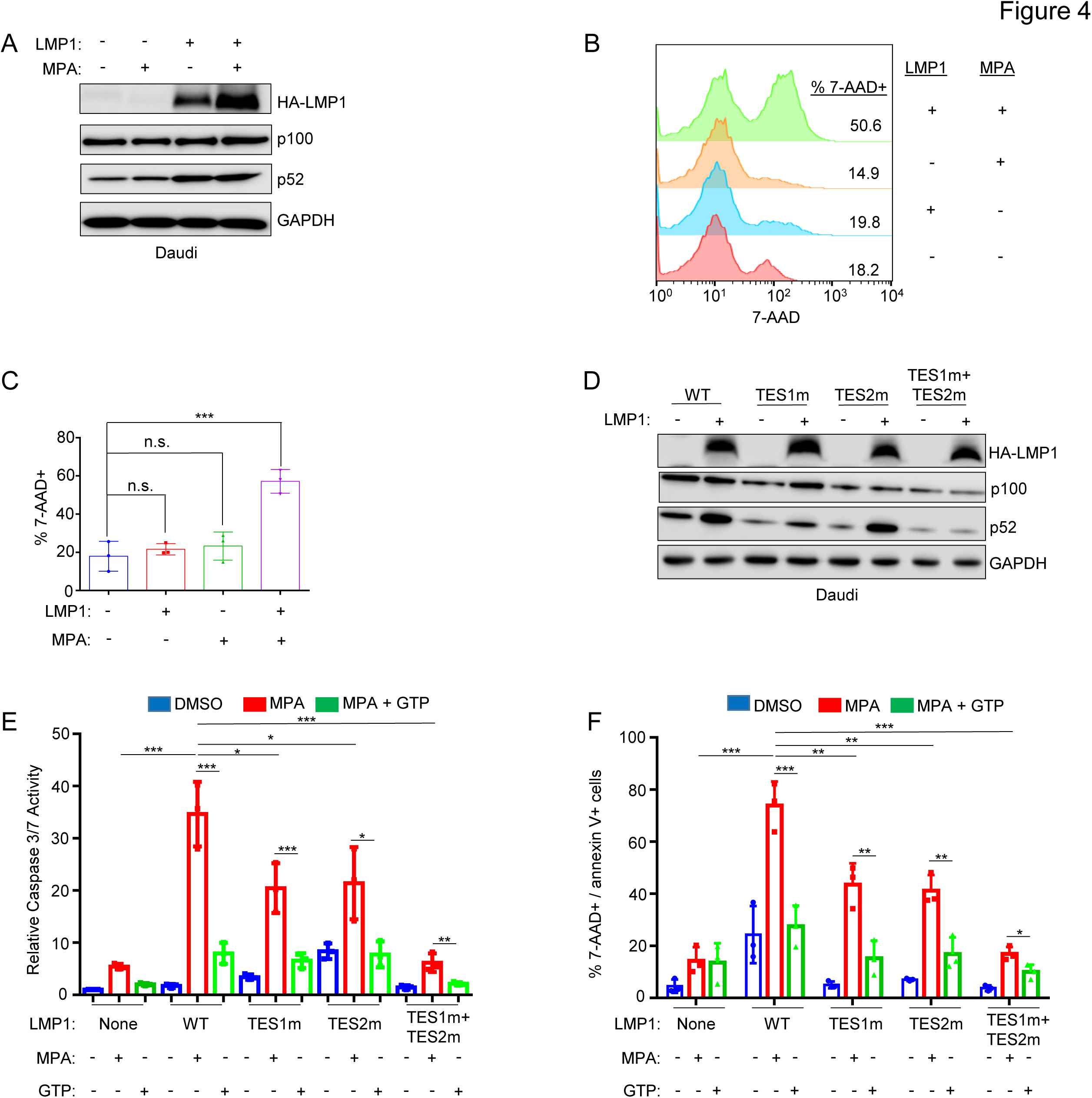
LMP1 expression sensitizes Burkitt cells to MPA-driven apoptosis in a partially GTP dependent manner. (A) MPA does not impair conditional LMP1 expression or signaling. WCL of Daudi cells induced for LMP1 and/or treated with MPA 1 μM for 24 hours, as indicated. (B) Analysis of LMP1 effects on MPA-driven Burkitt cell death. FACS analysis of 7-AAD uptake by Daudi cells mock induced or induced for LMP1 and then treated with DMSO vs MPA for 96 hours. Shown at right are the % 7-AAD+ cells under each condition. (C) Mean ± SD percentages of 7-AAD cells from n=3 replicates as in (B). (D) Validation of Daudi conditional wildtype vs mutant LMP1 expression. Immunoblot analysis of WCL of Daudi cells induced for wildtype (WT), TES1m, TES2m or TES1+TES2m LMP1 for 24 hours. (E) Analysis of TES1 vs TES2 effects on sensitization to MPA-induced Burkitt apoptosis. Relative caspase 3/7 activities in Daudi cells mock induced or induced for WT or the indicated LMP1 mutant for 24 hours and then treated with 1µM MPA ± 100mM GTP rescue for 96 hours. Caspase 3/7 levels were measured by Caspase-3/7 Glo assay, and values in mock induced and DMSO treated cells were set to 1. (F) Analysis of TES1 vs TES2 effects on sensitization to MPA-induced Burkitt apoptosis. Mean ± SD percentages of double 7-AAD+/annexin V+ (apoptotic cells) from n=3 replicates of Daudi cells treated as in (E). Immunoblots are representative of n=3 replicates. *P<0.05, **P<0.01, ***P<0.005.

To investigate whether signaling from the TES1 and/or TES2 domains were responsible for LMP1-driven IMPDH dependency, we conditionally expressed wildtype LMP1 or point mutants abrogated for signaling by TES1 (TES1 mutant, TES1m), TES2 (TES2m), or both (TES1m + TES2m)^62–64^. We validated that the TES1 point mutation abrogated non-canonical NF-κB pathway p100 to p52 processing as expected (**Figure 4D**). Intriguingly, MPA more highly induced apoptosis in cells with wildtype than with either TES1m or TES2m LMP1 expression (**Figure 4E-F, S6**). By contrast, expression of the TES1m+TES2m double LMP1 point mutant did not sensitize Daudi cells to MPA-driven apoptosis (**Figure 4E-F, S6**). These results indicate that signaling by either TES1 or TES2, rather LMP1 expression itself, contribute to dependence on IMPDH for survival. Similar results were obtained in EBV-negative BL-41 Burkitt cells, where MPA treatment again induced significantly higher apoptosis levels in cells with wildtype than TES1 or TES2 mutant LMP1 expression. These phenotypes were partially rescuable by GTP supplementation (**Figure S7-8**), suggesting that TES1 and TES2 signaling each increase GTP biosynthesis to meet increased demand.

### TES1 and TES2 each remodel B-cell metabolism

To gain further insights into TES1 and TES2 metabolism remodeling roles including at the nucleotide biosynthesis level, we performed LC/MS profiling of Burkitt cells at 24 hours post LMP1 TES1m versus TES2m expression. These samples were run in the same biological replicates as ones from cells mock induced or induced for wildtype LMP1 expression (**Fig. 1**) to facilitate cross-comparison and to limit batch effects. Interestingly, expression of TES1 mutant LMP1 (which signals from TES2) only mildly upregulated XMP and downregulated GTP abundance, relative to levels observed in mock induced Daudi cells (**Figure 5A**). TES2 mutant LMP1 (which signals from TES1) instead upregulated XMP abundance by ∼2-fold and significantly increased GMP levels (**Figure 5B**). TES2 mutant LMP1 likewise upregulated Akata XMP levels by ∼4-fold, whereas levels were again only modestly increased by TES1 mutant LMP1 (**Fig. S9A-B**). These data suggest that TES1 signaling may more strongly induce IMPDH activity and/or that TES2 signaling more strongly increases XMP demand.

**Figure 5.**
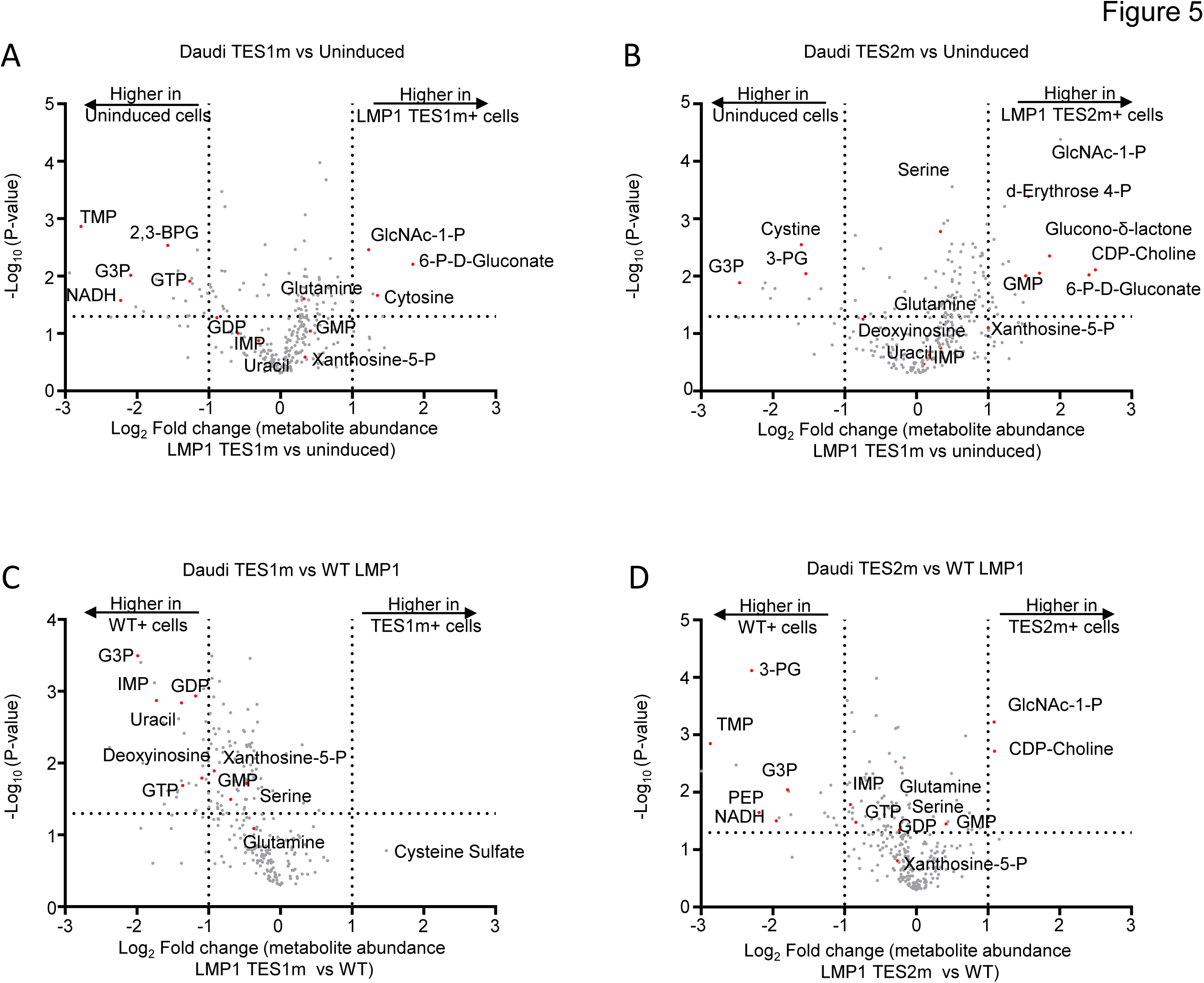
Effects of LMP1 TES1 versus TES2 signaling on Daudi Burkitt metabolome remodeling. (A) Volcano plot of LC-MS metabolomic analysis of n=3 replicates of Daudi cells mock induced or doxycycline induced for LMP1 TES1m expression for 24 hours. Metabolites with higher abundance in LMP1 TES1+ cells have positive fold change values, whereas those higher in mock induced cells have negative fold change values. Selected metabolites are highlighted by red circles and annotated. (B) Volcano plot of LC-MS metabolomic analysis of n=3 replicates of Daudi cells mock induced or doxycycline induced for LMP1 TES2m expression for 24 hours, with selected metabolites highlighted as in (A). (C) Volcano plot of LC-MS metabolomic analysis of n=3 replicates of Daudi cells doxycycline induced for TES1m vs WT LMP1 expression for 24 hours, with selected metabolites highlighted. Replicates for this cross-comparison were induced side by side, prepared for and analyzed by LC-MS together on the same day to minimize batch effects. (D) Volcano plot of LC-MS metabolomic analysis of n=3 replicates of Daudi cells doxycycline induced for TES2m vs WT LMP1 expression for 24 hours, with selected metabolites highlighted. Replicates for this cross-comparison were induced side by side, prepared for and analyzed by LC-MS together on the same day to minimize batch effects.

We next cross-compared metabolite abundances in Daudi cells expressing TES1 mutant versus wildtype LMP1. Wildtype LMP1 induced significantly higher levels of the GMP/AMP precursor inosine monophosphate (IMP), GDP and GTP than TES1 mutant LMP1. Deoxyinosine was also significantly higher in both Daudi and Akata expressing wildtype than TES1 mutant LMP1 (**Figure 5C, S9C**), further implicating TES1 signaling in nucleotide metabolism remodeling. By contrast, GDP or GMP levels were not significantly different in cells expressing TES2 mutant versus wildtype LMP1 (**Figure 5D, S9D**). Taken together, these results indicate that TES1 signaling may more strongly induce purine synthesis to support of GTP abundance.

### IMPDH metabolism crosstalk with EBV epigenome heterochromatin

We observed that MPA treatment induced Burkitt cell homotypic adhesion, in which large clusters or MPA-treated MUTU I or Daudi cells were evident (**Figure 6A**). Since LMP1 induces B-cell homotypic aggregation likely by upregulating plasma membrane L-selectin and other adhesion molecules^63,65^, we investigated whether MPA de-repressed Burkitt LMP1 expression. Immunoblot analysis demonstrated LMP1 expression in MPA-treated MUTU I, Akata, and Rael Burkitt cells, which could be suppressed by GTP supplementation (**Figure 6B**). Of note, Daudi LMP1 migrated at a lower molecular weight than LMP1 expressed within other MPA-treated Burkitt cells or GM15892 LCLs (**Figure 6B**). Since Daudi harbor an EBV genomic deletion that removes *EBNA2*, MPA effects on LMP1 were not indicative of a switch to the latency III program, and EBNA2 was not de-repressed in MUTU I, which have intact EBV genomes (**Supplemental Fig S10A**). LMP2A was also not de-repressed by MPA in either Daudi or MUTU I (**Supplemental Fig S10A**).

**Figure 6.**
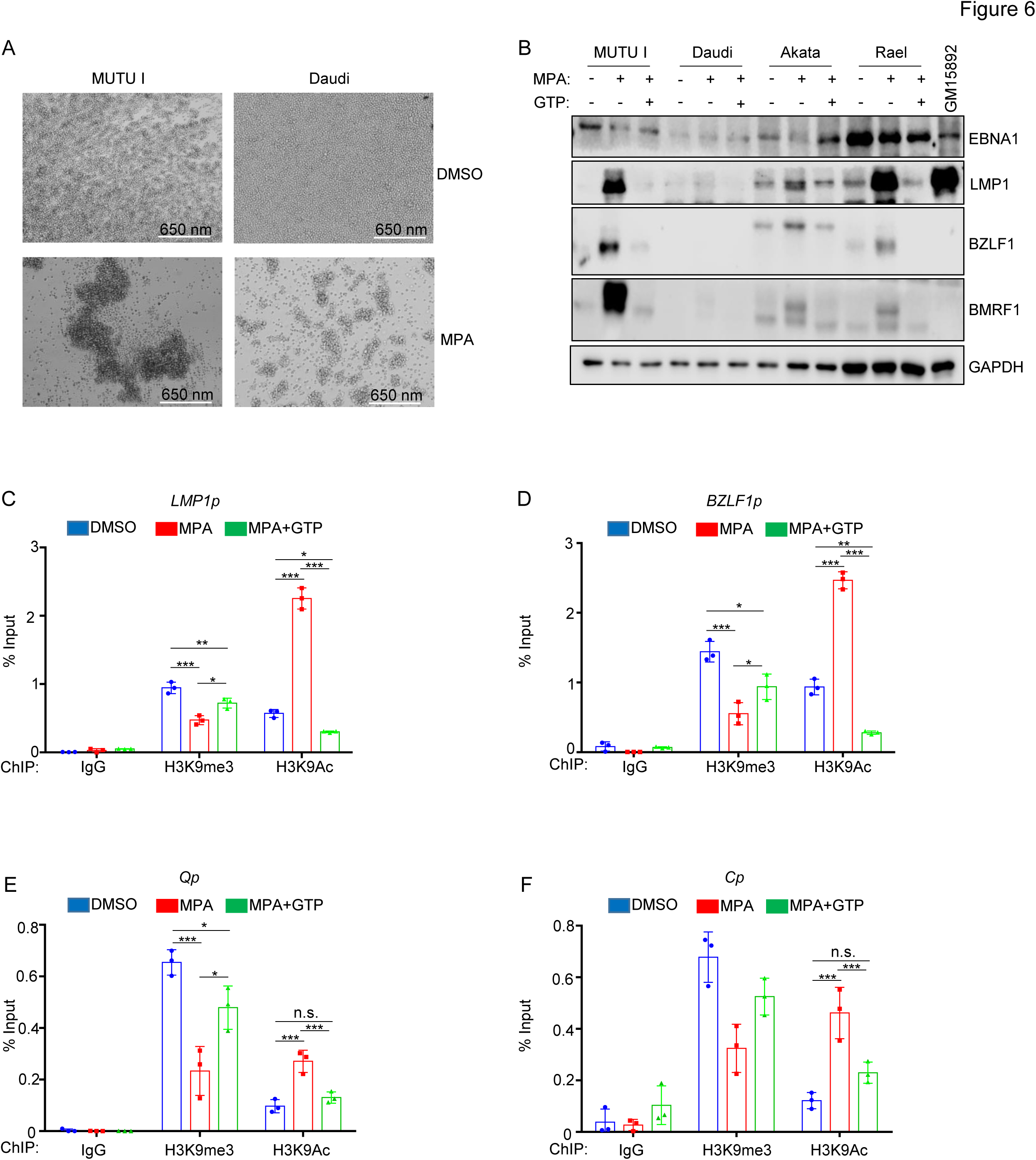
MPA derepresses lytic gene expression including LMP1 in latency I Burkitt cells in a GTP dependent manner. (A) Analysis of MPA effects on latency I Burkitt cell homotypic adhesion (cell clumping), a phenotype that typically correlates with LMP1 expression. Representative brightfield microscopy images of MUTU I and Daudi cells treated with DMSO or the indicated MPA concentration for 72 hours indicating MPA-induced homotypic adhesion. (B) Analysis of MPA effects on Burkitt LMP1, EBNA1, immediate early BZLF1 and early lytic BMRF1 expression. Immunoblot analysis of WCL from the indicated Burkitt cells mock treated or treated with 1µM MPA ± 100µM GTP for 24 hours. Shown in the rightmost lane are WCL from latency III GM15892 LCLs for cross comparison. MPA effects on LMP1 and on lytic gene expression were largely reversed by GTP supplementation. Blots are representative of n=3 replicates. (C) Mean ± SD percentage of input values from Daudi Burkitt cell chromatin immunoprecipitation (ChIP) with qPCR analysis, using the indicated control IgG, anti-H3K9me3 or anti-H3K9Ac antibodies, as indicated and with primers specific for the LMP1 promoter region (LMP1p). (D) Mean ± SD percentage of input ChIP-qPCR values as in (C) using primers specific for the immediate early BZLF1 promoter (BZLF1p) region. (E) Mean ± SD percentage of input ChIP-qPCR values as in (C) using primers specific for the latency I Q promoter (Qp) region. (F) Mean ± SD percentage of input ChIP-qPCR values as in (C) using primers specific for the latency III C promoter (Cp) region. *, P<0.05; **P<0.01, ***P<0.005, ns=non-significant.

Since LMP1 can be expressed as a latency gene or by the EBV lytic cycle^66,67^, we tested whether MPA de-repressed other EBV lytic cycle antigens. Immunoblot analysis demonstrated that MPA de-repressed immediate early BZLF1 and early BMRF1 in MUTU I, Akata and Rael. By contrast, neither LMP1 or lytic cycle antigens were appreciably de-repressed by MPA in Daudi cells. MPA effects on lytic cycle and LMP1 expression were rescuable by GTP supplementation (**Figure 6B**). These data are consistent with a model in which MPA de-represses LMP1 through Burkitt cell lytic reactivation.

To gain insights into MPA effects on the EBV epigenome, we performed chromatin immunoprecipitation (ChIP) and qPCR analysis. MPA reduced repressive histone 3 lysine 9 trimethyl (H3K9me3) marks and increased activating H3K9 acetyl (H3K9ac) marks at both the LMP1 and BZLF1 promoters. MPA also increased repressive histone 3 lysine 27 trimethyl (H3K27me3) but did not substantially alter H3K27 acetyl (H3K27ac) mark abundance at the LMP1 promoter (Figure S10B). MPA epigenetic effects were rescuable by GTP supplementation (**Figures 6C-D**). MPA induced similar epigenetic remodeling at the EBV genomic Q promoter, active in latency I, and at the latency III C promoter, repressed in latency I (**Figures 6E-F**). These results indicate that IMPDH activity broadly supports the latency I EBV epigenome rather than exerting specific effects at the LMP1 promoter. Taken together, our observations suggest that latency III B-cells depend upon LMP1-driven *de-novo* guanylate production for survival, whereas IMPDH1/2 activity instead maintains latency I Burkitt GTP levels for proliferation and latency maintenance (**Figure 7**).

**Figure 7.**
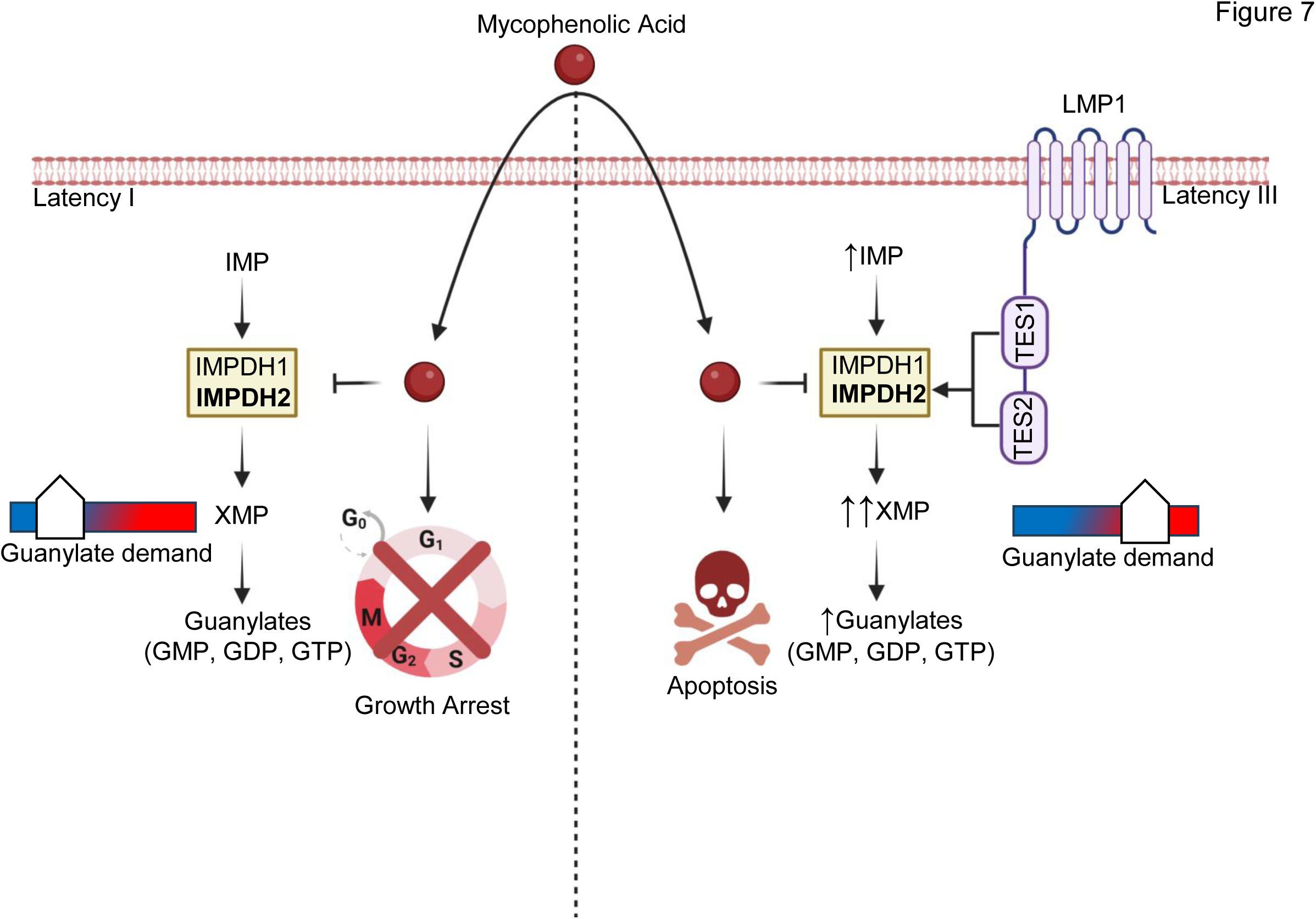
Schematic model of IMPDH1/2 inhibition effects on LMP1-negative latency I versus LMP1-positive latency III B-cell growth and survival. In Latency I, IMPDH1 and 2 each contribute to production of XMP and downstream guanylates to support demand. IMPDH1/2 inhibition by MPA triggers Burkitt growth arrest and de-represses EBV lytic antigens. In Latency III, LMP1 activated IMPDH2 predominantly supports XMP production and guanylate synthesis, sensitizing LMP1-expressing cells to MPA-driven apoptosis.

## Discussion

Metabolism remodeling is a cancer hallmark, yet much has remained unknown about how the key EBV oncogene LMP1 remodels host B-cell oncometabolism pathways. Here, LC/MS profiling highlighted that LMP1 significantly upregulates nucleotide metabolism pathways, in particular purine metabolism. While LMP1 did not alter IMPDH1 or IMPDH2 protein levels, it significantly upregulated levels of the IMPDH product XMP. While IMPDH blockade triggered Burkitt growth arrest, it instead triggered apoptosis in latency III Burkitt cells or LCLs, and LMP1 expression itself was sufficient to create dependence on IMPDH activity for survival. IMPDH2 played a larger role in LCL apoptosis evasion, though IMPDH1 also contributed. TES1 and TES2 signaling additively contributed to dependence on IMPDH activity for B-cell survival.

IMPDH regulates flux at the branch point between adenine and guanine nucleotide production and is the rate limiting step in guanine nucleotide biosynthesis^68^. A future objective will be to identify why LMP1 expression creates a metabolic dependency on IMPDH metabolism and GTP biosynthesis for B-cell survival, even in the context of its conditional Burkitt expression. Whereas lymphocytes typically rely on *de novo* biosynthesis to meet GTP demand, MPA does not induce B-cell apoptosis in most contexts, including of primary human peripheral blood or tonsil naïve, memory B-cells or plasmablasts^69^. Interestingly, despite the similarity between LMP1 and CD40 signaling pathways^8,9^, MPA also does not trigger apoptosis of primary human B-cells stimulated for 24 hours by CD40-ligand together with anti-IgM cross-linking and interleukin-21^69^. We therefore hypothesize that LMP1 signaling pathways not shared with CD40 markedly increase GTP demand, necessitating increased IMPDH2 and to a lesser extent IMPDH1 activity for B-cell survival.

Consistent with a recent report that EBNA2 induces IMPDH2^45,58–60^, we did not find evidence for induction of IMPDH1 or IMPDH2 mRNA or protein expression by LMP1. Since LMP1 expression instead increased IMPDH product XMP levels, our results suggest that LMP1 instead increases IMPDH activity, likely through post-translational modification. For instance, accumulation of the IMPDH precursor IMP drives formation of filamentous IMPDH structures termed cytoophidium, which can play key roles in maintaining GTP pools^70–73^. Cytoophidium reduce allosteric IMPDH1 and IMPDH2 inhibition by GTP and ATP^70,71,74,75^. IMPDH filaments are evident in proliferating germinal center B cells, presumably to meet elevated nucleotide demand in the highly-replicative dark zone^76^. However, we have not been able to obtain evidence for increased cytoophidium formation in LMP1+ cells. IMPDH1 can be phosphorylated at three sites, including by protein kinase Cα (PKC)^77^. Since LMP1 TES1/CTAR1 signaling activates PKC family members^30,78^, it will be of interest to determine whether they may mediate LMP1 effects on IMPDH. Alternatively, we observed increased ribose-5-phosphate and IMP abundance with LMP1 expression, and it remains possible that increased precursor levels drive flux through IMPDH to promote nucleotide biosynthesis.

Vertebrates encode two IMPDH enzymes, IMPDH1 and IMPDH2. Point mutations in each are linked to distinct human diseases^68^. Our CRIPSR analysis found that both IMPDH1 and IMPDH2 were important for latency I Burkitt B-cell proliferation, but IMPDH2 was more important for LCL survival than IMPDH1. Whereas IMPDH1 is widely expressed across most tissues, IMPDH2 is more abundantly expressed in activated lymphocytes^69,79^ and is nearly 5-fold more susceptible to MPA inhibition than IMPDH1^80^. We therefore speculate that LMP1 expression necessitates upregulation of IMPDH2 activity to meet the elevated GTP demand, in the absence of which apoptosis is triggered.

What then triggers apoptosis upon IMPDH inhibition of LMP1+ cells? p53 activation drives apoptosis in response to GTP depletion in renal cells^81^, although EBNA3C attenuates p53 function in latency III^82,83^. Nonetheless, LCLs have wildtype p53, and increases in p53 abundance such as by inhibition of the p53-targeting E3-ubiquitin ligase MDM2 can overcome EBNA3 restraint^84^. It will therefore be of interest to test whether CRISPR or chemical antagonism of p53 blocks MPA-driven apoptosis in LCLs or in LMP1+ Burkitt cells. It remains plausible that changes in GTP abundance could trigger a GTP-dependent apoptosis pathway^85^ or alter GTP-dependent trafficking of key dependency factors. For instance, we previously identified that LMP1 itself requires trafficking by the small GTPase Rab13 for effects on target gene regulation^44^.

LMP1 is expressed in a subset of epithelial tumors, in particular nasopharyngeal carcinoma (NPC)^86–88^. While specific NPC IMPDH roles reman unstudied, it is notable that IMPDH2 is expressed in NPC cell lines and in tumor tissues^89^. Interestingly, elevated IMPDH2 expression can serve as an independent prognostic biomarker for poor NPC prognosis in patients with localized or advanced metastatic disease^89^. While this study did not specifically examine LMP1 expression in NPC tumor tissues, it would be of interest to know whether cases with elevated IMPDH2 levels also exhibited abundant LMP1 expression. Relatedly, an important future objective will be to identify how LMP1 expression remodels purine nucleotide biosynthesis pathways, including GTP, in epithelial cells including in the nasopharyngeal and gastric carcinoma contexts. It will also be of interest to determine whether LMP1 expression creates epithelial cell metabolic dependency on IMPDH1 and/or 2, or whether this phenotype is specific to B-cells.

IMPDH2 is over-expressed in a range of cancers including glioblastoma, where its elevated levels drive increased synthesis of rRNA and tRNA, stabilization of the GTP-binding nucleolar protein nucleostemin and nucleolar hypertrophy^90^. Increased nucleolar size was observed by two days post-EBV infection of primary human B-cells, which reached peak size at day 4 post-infection, a timepoint where there are highly elevated levels of EBNA2 and MYC. MPA IMPDH inhibition reduced EBV-driven nucleolar hypertrophy and depleted nucleostemin at day 4 post-EBV infection^60^. Since LMP1 begins to be induced at the protein level by this timepoint^60^, it will be of interest to determine whether TES1 or TES2 signaling increases IMPDH2 activity and XMP abundance at this early timepoint, as well as to define the earliest timepoint post-EBV infection at which LMP1 expression reaches the threshold at which infected cells become sensitized to MPA-driven apoptosis.

MPA inhibits the outgrowth of newly infected primary human B-cells, reduces size of EBV-transformed cord blood B-cell xenografts *in vivo* and increases survival of mice with EBV-transformed B-cell xenograft tumors^60,91^. Notably, MPA treatment greatly reduced LMP1 expression in surviving xenograft tumor cells^60^, which we speculate occurred as a result of strong pro-apoptotic selective pressure generated by IMPDH inhibition in cells that retained LMP1 expression. It will be of interest to define how LMP1 is silenced in this setting, presumably by EBV epigenomic changes potentially including DNA methylation or polycomb repressive complex II activity^92^. Likewise, the MPA prodrug mycophenolate mofetil (MMF) is commonly used as part of immunosuppressive regimens post-transplant or with autoimmunity. It will be of interest to define whether EBV-driven lymphoproliferative diseases that break through IMPDH antagonists in patient populations exhibit diminished LMP1 expression, and if so, whether synthetic lethal approaches can be devised to selectively target such adapted EBV-infected B-cells.

In summary, LMP1 expression remodels host B-cell nucleotide *de novo* guanine nucleotide metabolism. LMP1 increased abundance of the IMPDH product XMP and created metabolic dependency on IMDPH activity for B-cell survival. IMPDH2 played a more important role in LCL survival, though both IMPDH isoforms contributed to apoptosis blockade. Both TES1 and TES2 signaling supported guanine metabolism. Whereas IMPDH blockade triggered LCL apoptosis, it instead caused Burkitt growth arrest and LMP1 de-repression in the context of lytic reactivation. Our results further highlight IMPDH metabolic dependency as a rational therapeutic target for the treatment of EBV-driven immunoblastic lymphomas driven by LMP1.

## Acknowledgements.

This work was supported by an American Cancer Society Post-doctoral fellowship to E.M.B., by T32AI007245, and by R01DE033907, R01CA228700, R01AI164709, U01CA275301 and P01CA269043 to B.E.G.

## Materials and Methods

### Cell lines and culture

293T, Daudi and Jijoye were purchased from American Type Culture Collection. P3HR-1, Daudi, Kem I, and EBV-Akata^93^ were obtained from Elliott Kieff. GM11830, GM12878, GM13111, and GM12881 LCL were obtained from Coriell. Mutu I and Mutu III were obtained from Jeff Sample and Alan Rickinson. 2-2-3 LCLs with a conditional EBNA2-HT allele were obtained from Bo Zhao and Elliot Kieff. All B-cell lines were cultured in RPMI-1640 (Invitrogen) supplemented with 10% fetal bovine serum (FBS). 293T cells were cultured in DMEM with 10% FBS. All cell lines were incubated with 1% penicillin-streptomycin (Gibco) in a humidified incubator at 37ᵒC and 5% CO2. All cells were routinely confirmed to be mycoplasma-negative by Lonza MycoAlert assay (Lonza). For 2-2-3 cell assays, cells cultured in the presence of 4-hydroxy tamoxifen (4HT) exhibit latency III. The conditional EBNA2-HT localizes to the nucleus in the presence of 4HT, but upon 4HT withdrawal, it is sequestered in the cytosol. 2-2-3 LCLs were maintained in the presence of 1 μM 4HT. For 4HT removal, cells were washed five times with 4HT-free media. The first two washes included 30-minute incubations in 4HT free media. Cells were then seeded with or without 4HT for 48 hours before analysis.

**Primary B Cell Isolation and Culture.**

Platelet-depleted venous blood obtained from the Brigham & Women’s hospital blood bank were used for primary human B cell isolation, following our Institutional Review Board-approved protocol for discarded and de-identified samples. RosetteSep and EasySep negative isolation kits (Stemcell Technologies) were used sequentially to isolate CD19+ B cells by negative selection, with the following modifications made to the manufacturer’s protocols. For RosetteSep, 40 μL antibody mixture was added per mL of blood and before Lymphoprep density medium was underlayed, prior to centrifugation. For EasySep, 10 μL antibody mixture was added per mL of B cells, followed by 15 μL magnetic bead suspension per mL of B cells. After negative selection, the cells were washed twice with 1x PBS, counted, and seeded for EBV infection studies. Cells were cultured in RPMI-1640 (Invitrogen) supplemented with 10% FBS and penicillin-streptomycin in a humidified chamber at 37ᵒC and 5% CO2. Cells were cultured in RPMI-1640 supplemented with 10% FBS and penicillin-streptomycin in a humidified incubator at 37ᵒC and at 5% CO2.

### Antibodies and Reagents

Antibodies against the following proteins were used in this study: IMPDH1 (Cell Signaling Technology, #57068), IMPDH2 (Cell Signaling Technology, # 35914S), GAPDH (EMD Millipore, MAB374), LMP1 (Abcam, ab78113), LMP2A (Abcam, ab59028), EBNA2 PE2 (a gift from Fred Wang), DDX1 (Bethyl, A300-521A-M), Myc (Santa Cruz Biotechnology, SC-40), p100/p52 (EMD Milipore, 05-361), TRAF1 (Cell Signaling Biotechnology, #4715S), HA tag antibody (Cell Signaling Technology, # 3724), Fas-APC (Biolegend, 305612), ICAM-1-PE (BD Bioscience, 555511), Caspase 3 (Cell Signaling Technology, #9662), EBNA1 (a gift from Jaap Middledorp). The following reagents were utilized in this study at the indicated concentration unless otherwise noted: DMSO (Fisher, BPBP231-100), mycophenolic acid (Selleckchem Cat#S2487, 1 μM), doxycycline hyclate (Sigma, D9891-1G, 250 ng / mL), GTP (Roche, 10106399001, 100 μM).

**B95.8 EBV Preparation and B-cell Infection**

B95-8 EBV stocks were prepared from B95-8 producer cells as previously described^94,95^ and stored at -80 degrees C. Infectious titer of freshly thawed EBV was determined by primary B-cell transformation assay. Freshly isolated, deidentified, discarded CD19+ peripheral blood B cells were seeded in RPMI1640 with 10% FBS at a concentration of one million cells/mL for infection studies at an EBV multiplicity of infection of 0.1.

### Metabolite Extraction and Metabolomic Analysis

Metabolites were extracted according to Asara lab published protocols^96^. Twenty-four hours after doxycycline (250 ng / mL, Sigma Cat#D9891-1G) addition, uninduced and LMP1-induced Burkitt cells were pelleted and resuspended in fresh RPMI/FBS. After two hours, metabolites were extracted from two million cells using 4 mL of -80 degrees C 80% methanol, made from HPLC grade water (Sigma, 270733-1L) and LC-MS-grade methanol (Fisher, A456-1). Cells were suspended in 80% methanol by vortexing and pipetting, and extraction was performed overnight in a 4 degree C room on dry ice to ensure metabolite stability. Cell debris was pelleted in a 15 mL conical at 14,000*g* for 5 minutes at 4 degrees C and supernatant was collected on dry ice. An additional round of extraction was performed using 0.5 mL of fresh 80% methanol (-80 degrees C). Cell debris were vigorously suspended using a combination of intense vortexing and pipetting with a p1000. A total of 4.5 mL of extracted metabolites were split into three 1.5 ml Eppendorf tubes and then dried using a Speedvac for ∼6-8 hours. Dried pellets were stored at -80 until resuspension in HPLC water and LC-MS/MS analysis. LC-MS/MS was performed as published^96^. Peak area integrated TIC were utilized for relative comparisons of metabolites between samples.

### ChIP and ChIP-qPCR

Five million cells were pelleted, washed using 1x PBS, and then fixed with 1% formaldehyde (Sigma, Cat#252549-100ML) in RPMI1640 for 20 minutes at room temperature. Cross-linking was quenched by adding 2.5M glycine at room temperature for 5 minutes. After two more 1x PBS washes, Cells were lysed in 1 mL of ChIP lysis buffer (50 mM Tris, 10 mM EDTA, 1% SDS) supplemented with 1x cOmplete, EDTA-free Protease Inhibitor Cocktail (Pierce). Lysates were divided up into 250 uL aliquots and then sonicated with Bioruptor (Diagenode) with 30s on, 30s off for 12 cycles. Shearing of chromatin was confirmed by electrophoresis through a 0.8% Agarose gel. Sonicated chromatin was diluted 1:10 with ChIP dilution buffer (1.2 mM EDTA, 16.7 mM Tris, 167 mM NaCl, 0.01% SDS, 1.1% Triton X-100) and incubated with antibodies of interest or control IgG (Cell Signaling Technology, #2729S) antibodies overnight at 4c. Antibodies used for ChIP were anti-H3K9me3 (Active Motif, 39062), anti-H3K9Ac (Cell Signaling Technology, 9649S) anti-H3K27ac (Abcam, ab4729), and anti-H3K27me3 (Active Motif, 39155). Protein-DNA complexes were precipitated with protein A/G magnetic beads added at the same time as the antibodies (Pierce, 88803). Magnetic beads were washed extensively (washed twice with a lower salt buffer (150 mM NaCl, 2 mM EDTA, 20 mM Tris, 0.1% SDS, 1% Triton X-100) and then a high-salt buffer (500 mM NaCl, 2 mM EDTA, 20 mM Tris, 0.1% SDS, 1% Triton X-100), and once with LiCl buffer (0.25 M LiCl, 1% NP-40, 1% sodium deoxycholate, 1 mM EDTA, 10 mM Tris) and finally TE buffer (10 mM Tris, 1 mM EDTA). Each wash was 1 mL volume. Buffers were removed by placing solutions on magnet for 1 minute. Chromatin was eluted in Elution buffer (100 mM NaHCO3, 1% SDS) and reverse cross-linked at 65 °C for 2 hours. QIAquick PCR purification kits were used to purify the immunoprecipitated DNA, followed by qPCR with PowerUp SYBR green PCR master mix on a CFX Connect Real-Time PCR Detection System (Bio-Rad). qPCR was performed using 1 μL of eluted DNA per reaction, 0.5 μM FP and 0.5 μM RP for a total volume of 10 μL per reaction. Delta Ct values normalized to the percentage of input DNA. ChIP-PCR was performed using the primers below:

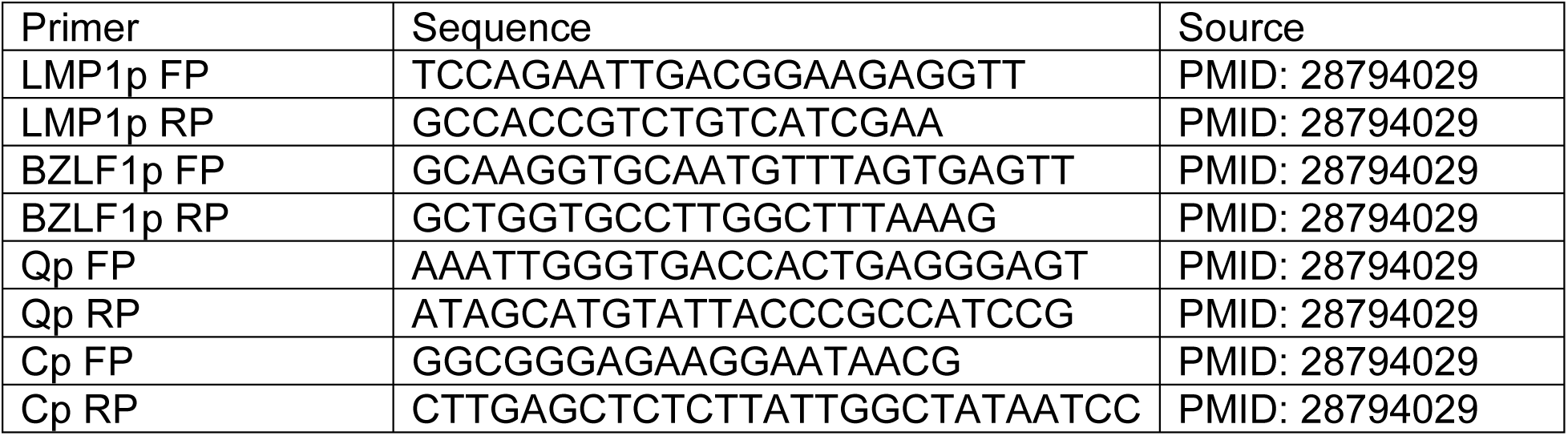

### Immunoblot analysis

Cells were lysed in Laemelli buffer (0.0625 M Tris Base, 0.07M SDS, 10% glycerol v/v, 5% 2-mercaptoethanol, .002% bromophenol blue) and sonicated at 4 degrees C for 10 seconds using a probe sonicator at the maximum setting. Lysates were centrifuged at 17,000 x g for 30 minutes and harvested supernatants were boiled for 10 minutes. SDS-PAGE was performed using 8 or 10% acrylamide gels and transferred onto nitrocellulose membranes for 90 minutes at 100 V at 4°C. Membranes were blocked for 1 hour at room temperature in 5% non-fat milk / 1x TBST. Blots for phosphorylated residues were blocked using 2% BSA instead of 5% non-fat dry milk in TBST. Blocked membranes were probed with primary antibodies (diluted in 1x TBST with 0.02% sodium azide at recommended manufacturer concentrations) overnight at 4°C, followed by a one hour incubation with respective HRP-conjugated secondary antibodies in 1x TBST. Blots were imaged on a LiCor Odyssey workstation.

### CRISPR/Cas9 mutagenesis

B-cell lines with stable Cas9 expression were established as described previously^97^. sgRNA constructs were generated as previously described^98^ using sgRNA sequences from the Broad Institute Avana or Brunello libraries. CRISPR editing was performed as previously described^99^. Briefly, lentiviruses encoding sgRNAs were generated by transient transfection of 293T cells with packaging plasmids pasPAX2 (Addgene, Plasmid #12260) and pCMV-VSV-G (Addgene, Plasmid #8454). and pLentiGuide-Puro (Addgene, Plasmid #52963) plasmids. P3HR-1, Daudi, Mutu I, GM12878, and GM13111 cells stably expressing Cas9 were transduced with the lentiviruses and selected with 3 μg/mL puromycin (Thermo Fisher, Cat#A1113803) for three days before replacement with antibiotic-free media. CRISPR editing was confirmed by immunoblotting 3 days post puromycin selection. For IMPDH1/IMPDH2 double knockout, in addition to the pLentiGuide Puro vector harboring IMPDH1 sgRNA a, second transduction was performed 3 days after the addition of puromycin using pLenti SpBsmBI sgRNA Hygro vector (Addgene, Plasmid # 62205) harboring the IMPDH2 sgRNA. Hygromycin (100 μg/mL, Thermo Fisher, Cat #10687010) selection was carried out two days post the second transduction. The sgRNAs used in this study were constructed using oligos based on the sequences below:

**Table S1:**
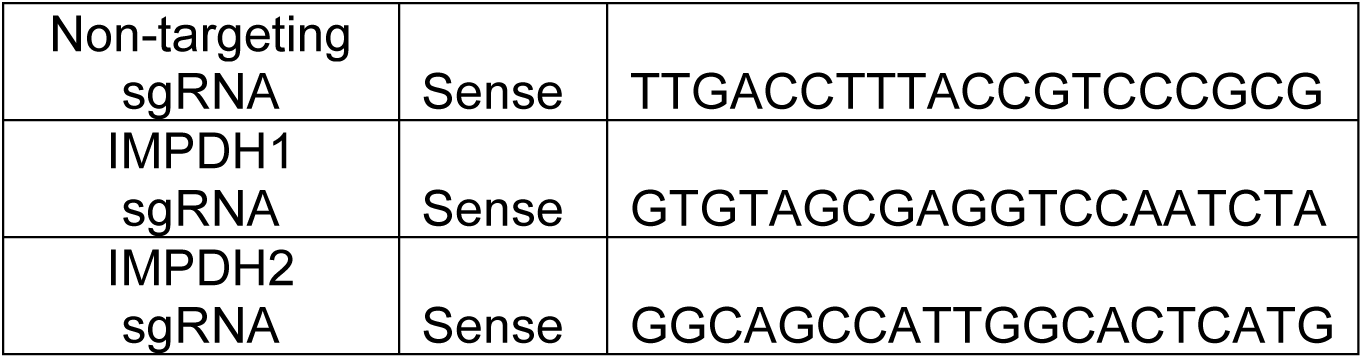
Table of sgRNAs used to knockout genes of interest in this study. sgRNAs used for CRISPR-Cas9-mediated knockout of the indicated genes are listed above.

### Flow cytometry

For Fas / ICAM-1 staining, cells were harvested and washed twice with 1x PBS supplemented with 2% FBS (Gibco). Cells were then incubated 45 minutes at room temperature with indicated antibodies at manufacturer concentrations. Cells were washed twice more after incubation to remove excess antibody and immediately analyzed on a flow cytometer. For 7-AAD (Thermo Fisher, Cat#A1310) viability assays, cells were harvested and washed twice with 1x PBS supplemented with 2% FBS (Gibco). Cells were then incubated with a 1 μg / mL 7-AAD solution in 1x PBS / 2% FBS for five minutes at room temperature, protected from light. Cells were then analyzed via flow cytometry. For Annexin V (Biolegend, Cat#640906) vs 7-AAD assays, cells were harvested and incubated with a solution containing: 2.5 μL of Annexin V FITC antibody, 2.5 μL of a 20 ng / mL solution of 7-AAD, and 100 uL of with 1x PBS supplemented with 2% FBS per sample. Cells were incubated in this Annexin V / 7-AAD solution for 20 minutes at room temperature, protected from light, before immediate analysis. For CFSE labeling and cell proliferation assays, CellTrace™ CFSE (Invitrogen, Cat#C34554) solution was prepared according to manufacturer’s instructions. 10^7 cells were resuspended and incubated in one mL of CellTrace™ working solution for ten minutes in a 37°C / 5% CO2 incubator, protected from light, with the cap of the vessel ajar. Five mL of RPMI 1640 with 10% FBS was added to the stained cells protected from light. Cells were incubated at room temperature for five minutes, protected from light, to remove free dye and prevent toxicity. Cells were then pelleted by centrifugation (300g x 5 minutes, room temperature) and resuspended in fresh RPMI with 10% FBS three times before the cells were seeded in complete media for proliferation analysis experiments at a concentration of 300,000 cells/mL. Labeled cells were analyzed by flow cytometry using a BD FACSCalibur instrument and analysis was performed with FlowJo V10.

### CellTiter-Glo

CellTiter-Glo viability assay (Promega) was performed according to the manufacturer’s protocol at the indicated time points. A total of 50 μL cells in PBS were used per assay according to manufacturer instruction.

### Software/data Presentation

Statistical analysis was assessed with Student’s t test using GraphPad Prism 7 software, where NS = not significant, p > 0.05; * p < 0.05 ** p < 0.01; *** p < 0.005. and graphs were made using GraphPad Prism 7. Schematic models were made using Biorender. Metabolic pathway analyses were performed using Metaboanalyst 6.0 platform^51^.

---

**Figure S1.**
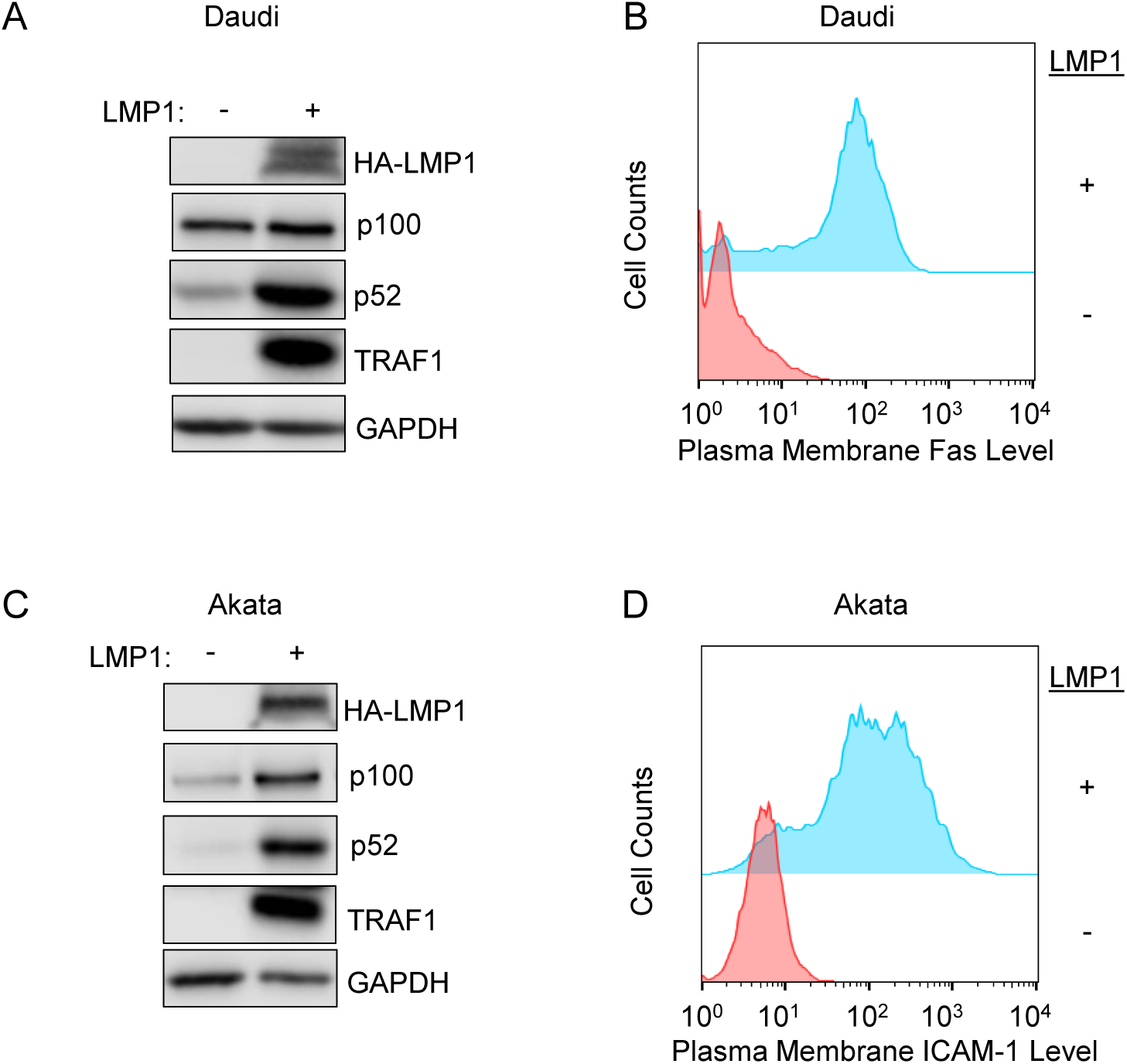
Validation of conditional LMP1 Burkitt cell models. (A) Immunoblot analysis of whole cell lysates (WCL) from Daudi cells mock induced or induced for LMP1 expression by doxycycline (250ng/mL) for 24 hours. LMP1 expression induced non-canonical NF-κB activity, as judged by processing of the p100 precursor into the p52 subunit, and induced expression of the well characterized LMP1/NF-κB target TRAF1. (B) FACS analysis of plasma membrane Fas abundance in Daudi cells mock induced or induced for LMP1 expression for 24 hours, as in (A). Conditional LMP1 expression highly induced expression of the well characterized LMP1 target Fas. (C) Immunoblot analysis of WCL from Akata cells mock induced or induced for LMP1 expression as in (A), indicating LMP1 induction of non-canonical NF-κB activity and LMP1 target TRAF1 expression. (D) FACS analysis of plasma membrane ICAM-1 abundance in Akata cells mock induced or induced for LMP1 expression as in (A). ICAM-1 rather than Fas was analyzed as basal Fas expression is aberrantly elevated in Akata cells.

**Figure S2.**
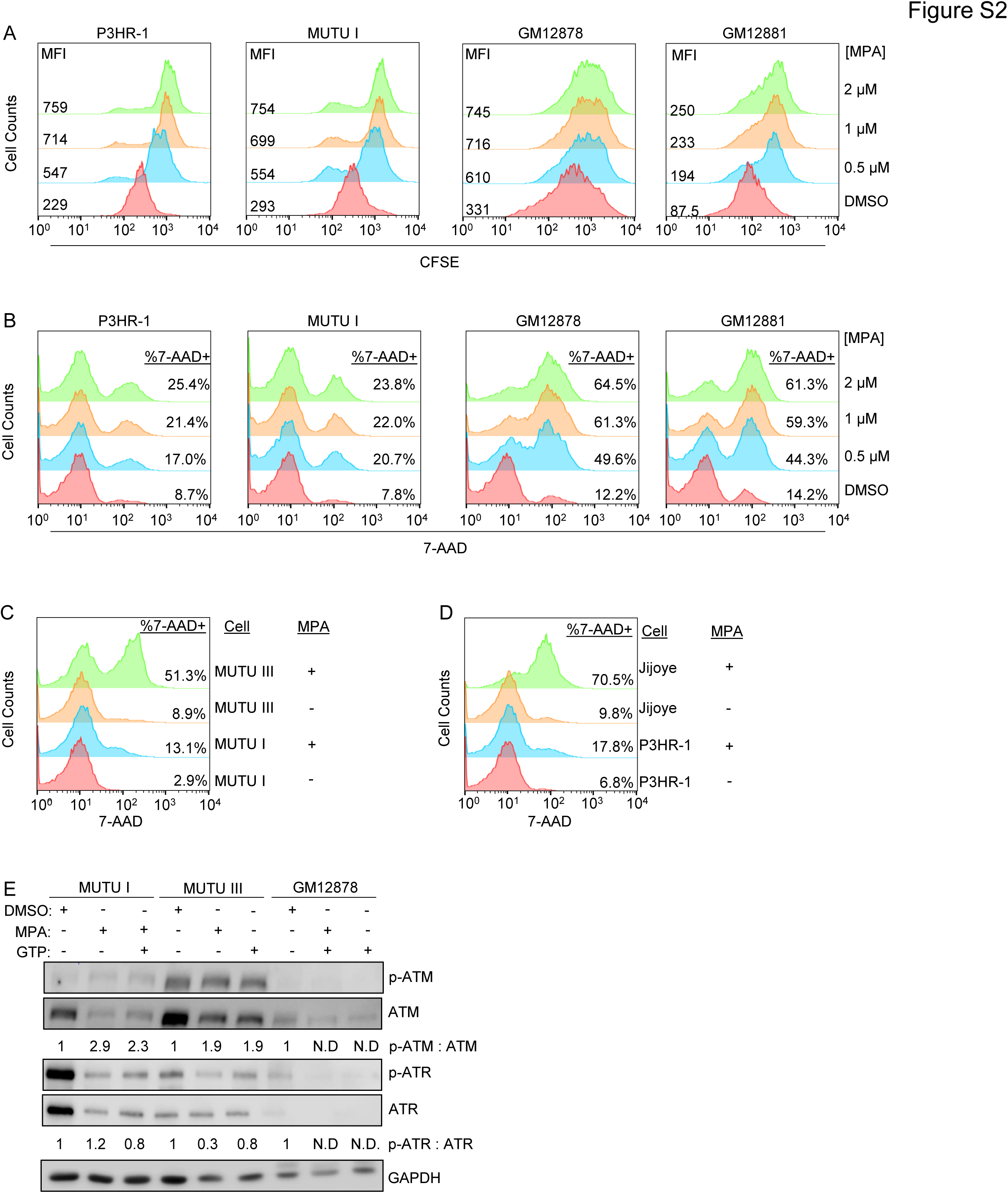
EBV latency state dependent MPA effects on proliferation and survival. (A) FACS analysis of CFSE levels following 96 hours of treatment with the indicated MPA dosages in the indicated cell lines. (B) FACS analysis of 7-AAD uptake following 48 hours of treatment of indicated cell lines with the indicated MPA dosages. (C) FACS analysis of MUTU I versus III 7-AAD uptake following 48 hours of treatment with the indicated MPA dosages. (D) FACS analysis of P3HR-1 versus Jijoye cell 7-AAD uptake following 48 hours of treatment of MUTUI vs III with the indicated MPA dosages. (E) Immunoblot analysis of WCL from MUTU I, III or GM12878 treated with DMSO or 1 μM MPA with or without 100 μM GTP rescue for 48 hours. Shown are the normalized phospho-ATM:ATM and phospho-ATR:ATR ratios calculated by densitometry analysis, with values in DMSO-treated cells set to 1. N.D., not determined.

**Figure S3.**
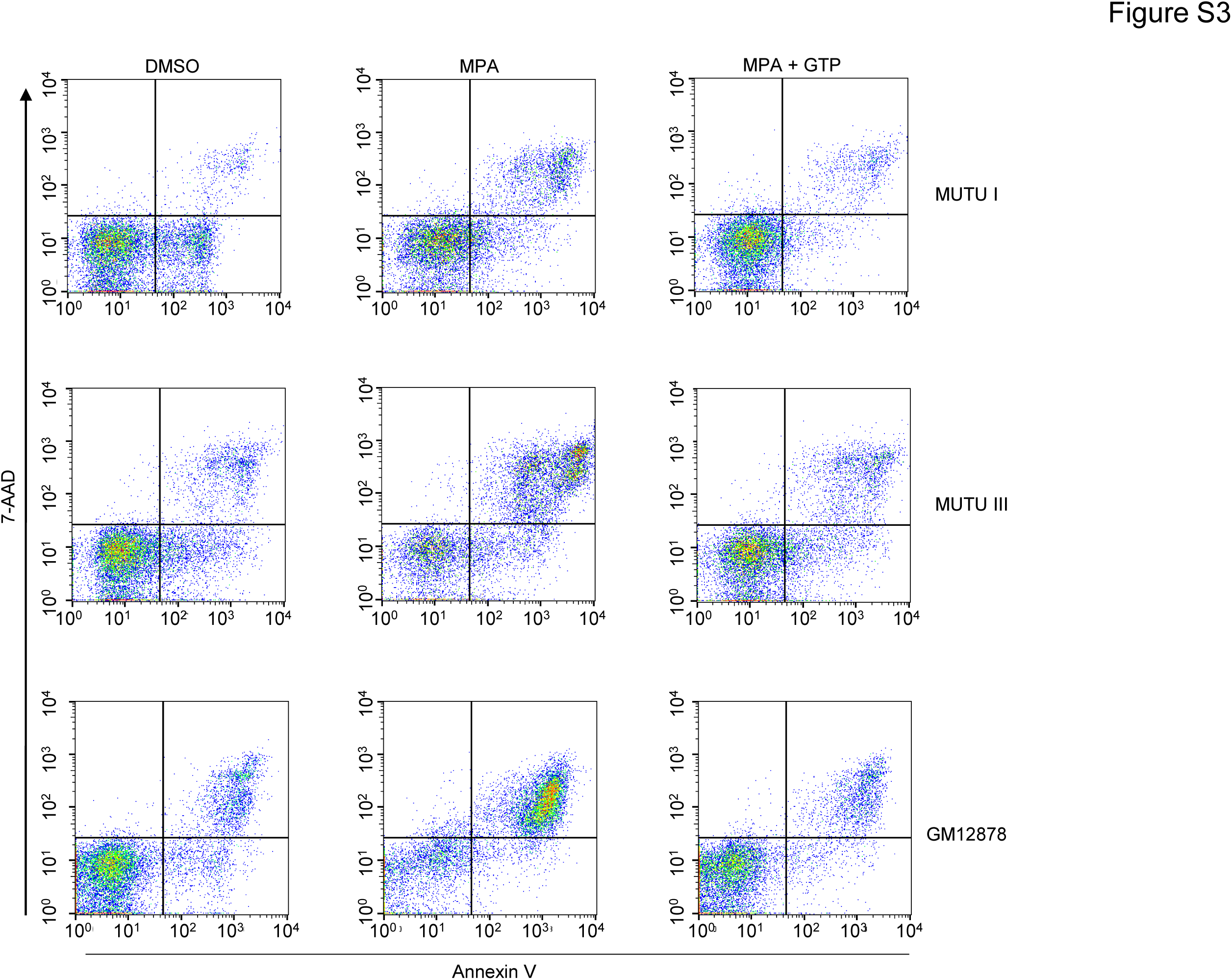
FACS analysis of MPA effects on GTP-dependent EBV+ B-cell viability in latency I versus III. Shown are representative FACS plots from n=3 replicates of the indicated cell lines treated with DMSO, 1 μM MPA or 1 μM MPA + 100 μM GTP for 48 hours, as shown in Figure 2F.

**Figure S4.**
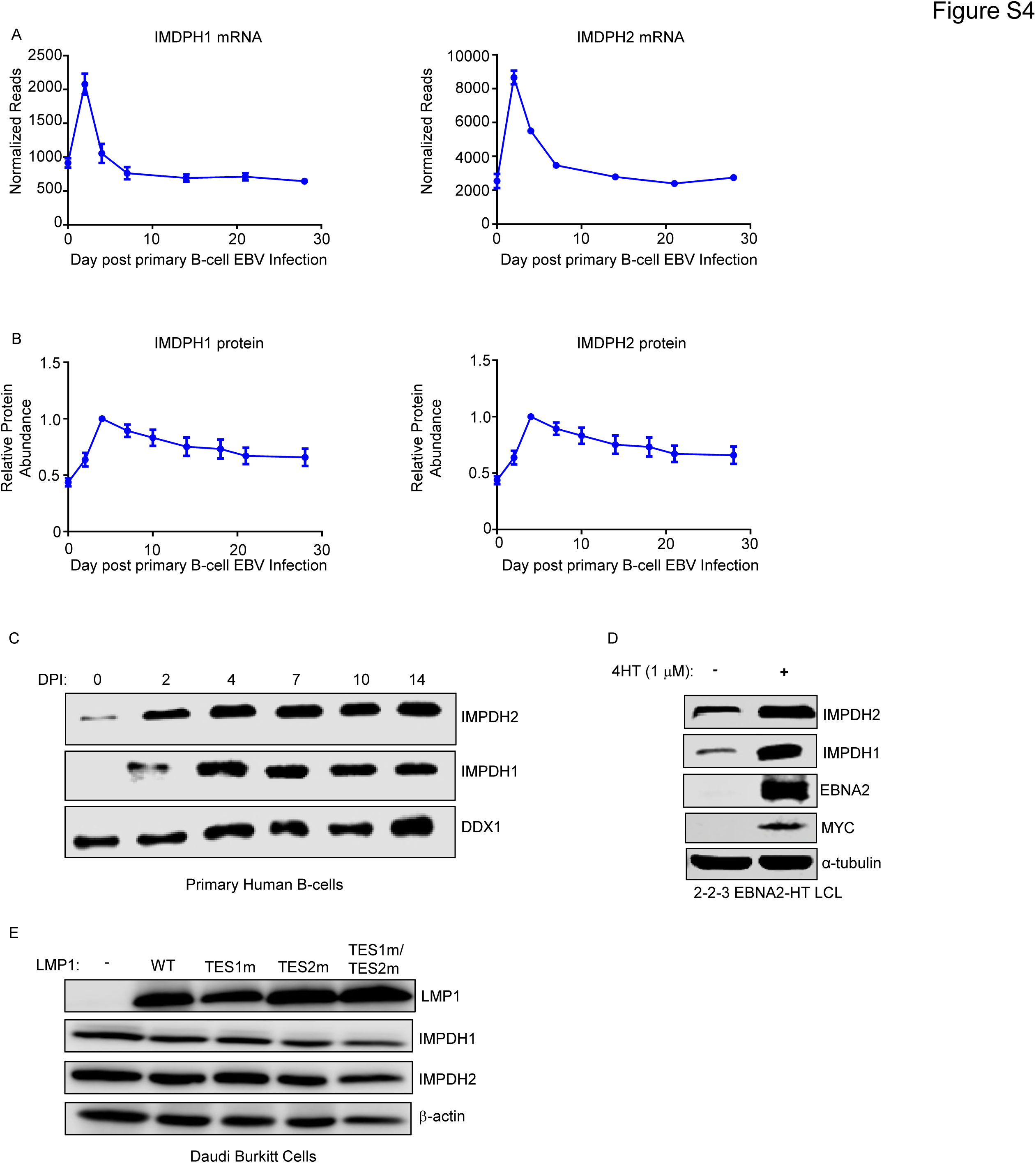
EBV latency gene effects on IMPDH1 and IMPDH2 expression. (A) Normalized *IMPDH1* (left) versus *IMPDH2* (right) reads from RNAseq analysis of primary human B cells at the indicated days post-infection by the EBV B95.8 strain. Shown are mean ± SD values from n=3 replicates^45,58–60^. (B) Relative IMPDH1 (left) versus IMPDH2 (right) protein abundances from tandem mass tagged multiplexed mass spectrometry proteomic analysis of primary human B-cells at the indicated days post-infection by the EBV B95.8 strain. Shown are mean ± SD values from n=4 replicates^45^. (C) Immunoblot analysis of WCL from primary human B-cells infected by B95.8 EBV at the indicated days post infection (DPI). DDX1 was used as a load control as its levels remain relatively unchanged between uninfected versus infected primary B-cells^45^. (D) Immunoblot analysis of WCL from 2-2-3 EBNA2-HT LCLs, which harbor a conditional EBNA2 allele that is fused to a modified estrogen receptor ligand binding domain, whose activity requires the presence of 4-hydroxytamoxifen (4-HT). WCL were generated at 48 hours post EBNA2 inactivation by 4-HT washout. (E) Immunoblot analysis of WCL from Daudi cells mock induced or induced for wildtype, TES1 point mutant (TES1m), TES2 point mutant (TES2m), or double TES1m/TES2m LMP1 for 24 hours by 250 ng/mL doxycycline. Immunoblots are representative of n=3 independent replicates

**Figure S5.**
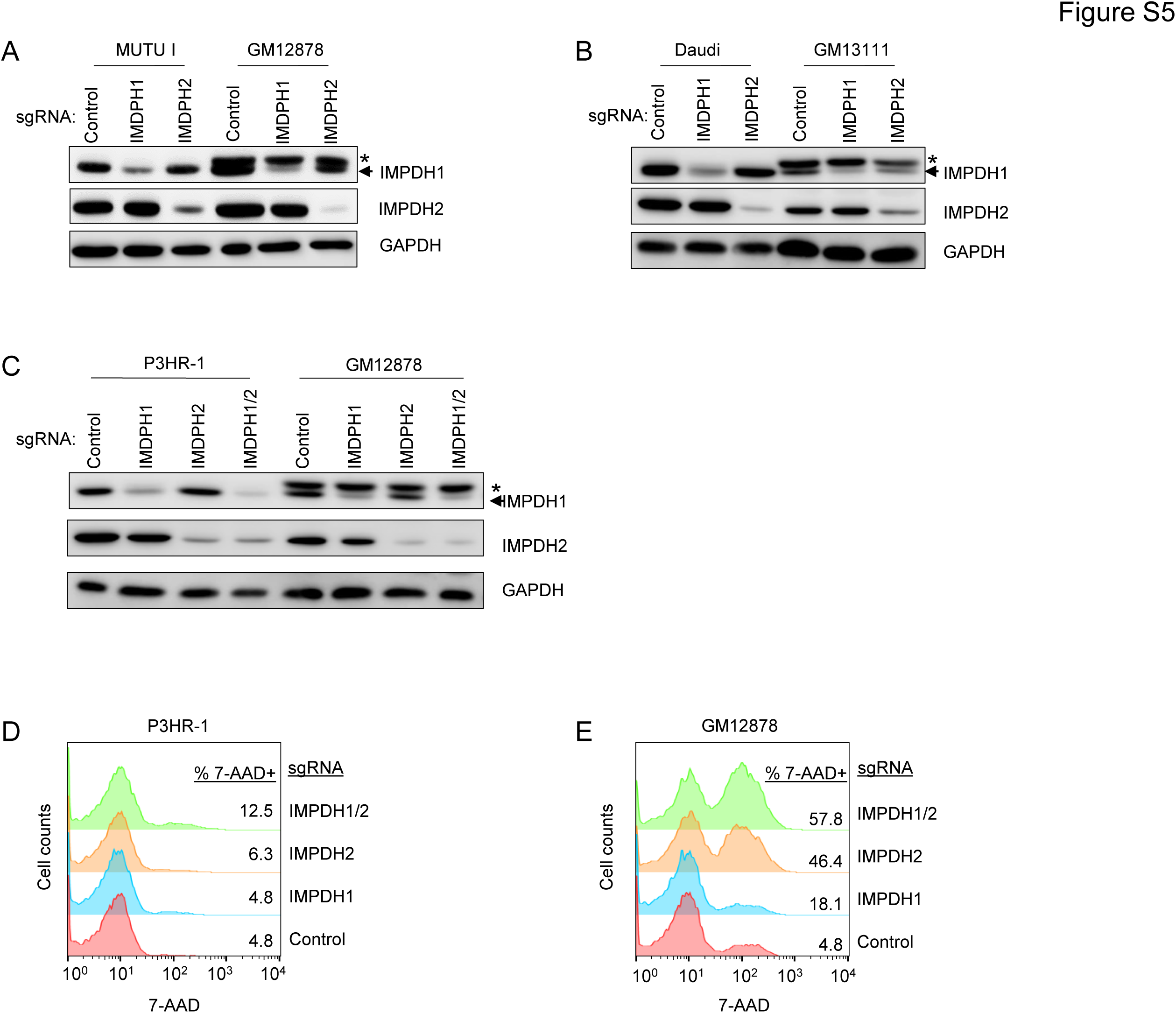
Genetic perturbation of IMPDHs impact growth and viability of lymphoblastoid cell lines. (A) Immunoblot analysis of IMPDH1 or IMPDH2 depletion. WCL from Cas9+ MUTU I or GM12878 expressing were subjected to immunoblot analysis at the indicated sgRNA at two days post-puromycin selection of successfully transduced cells. *=non-specific band. (B) Immunoblot analysis of IMPDH1 or IMPDH2 depletion in Daudi versus GM13111 WCL, as in (A). (C) Immunoblot analysis of IMPDH1 or IMPDH2 depletion in P3HR-1 versus GM12878 WCL, as in (A). (D) FACS analysis of 7-AAD uptake in Cas9+ P3HR-1 expressing the indicated sgRNA at 8 days post-puromycin selection of cells transduced by sgRNA expressing lentivirus. (E) FACS analysis of 7-AAD uptake in Cas9+ GM12878, as in (D). FACS plots and immunoblots are representative of n=3 independent replicates.

**Figure S6.**
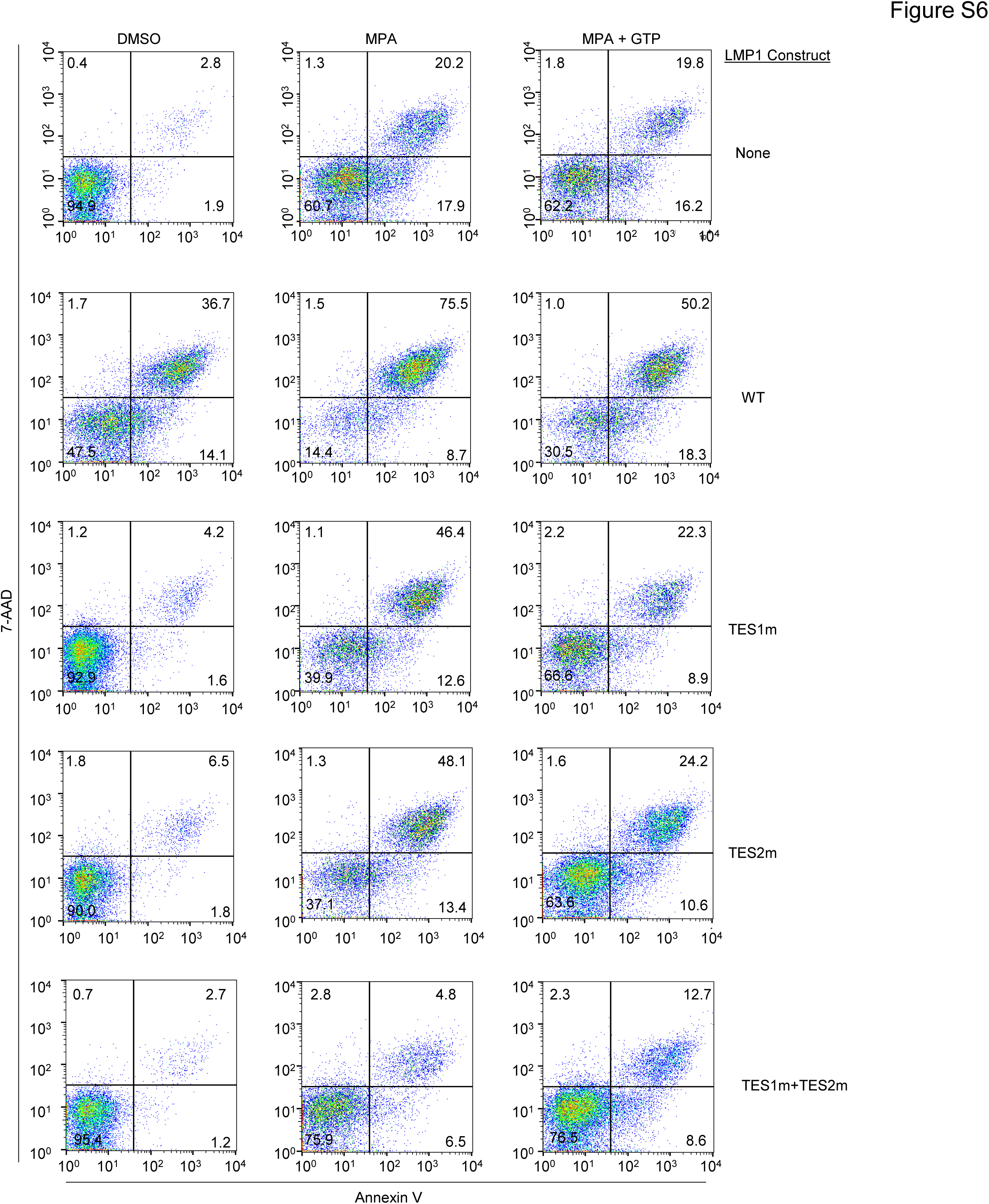
LMP1 expression sensitizes Daudi cells to MPA-driven apoptosis in a partially GTP dependent manner. Representative FACS plots from n=3 replicates of Daudi cells mock induced or induced for WT, TES1m, TES2m or TES1m+TES2m LMP1 expression for 24 hours and then treated with 1μM MPA ± 100μM GTP for 96 hours as indicated and as in Figure 4. Shown are FACS analysis of 7-AAD uptake versus Annexin V positivity.

**Figure S7.**
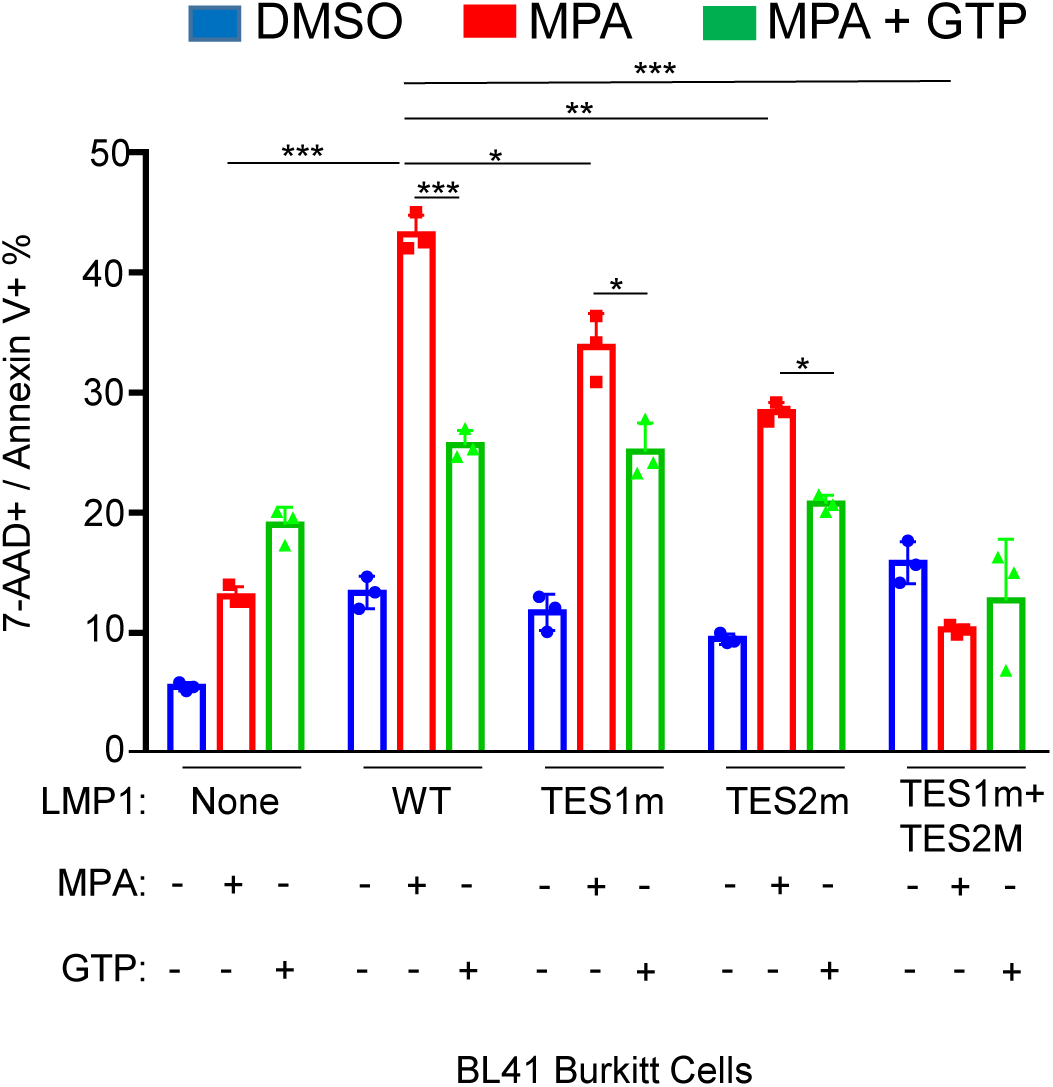
LMP1 TES1 and TES2 signaling sensitize EBV-negative BL41 Burkitt cells to MPA-driven apoptosis in a partially GTP dependent manner. Mean ± SD percentages of double 7-AAD+/annexin V+ (apoptotic cells) from n=3 replicates of BL41 cells mock induced or induced for the indicated LMP1 construct for 24 hours and then treated with DMSO or 1μM MPA ±100 μM GTP for 96 hours, as indicated. *P<0.05, **P<0.01, ***P<0.005.

**Figure S8.**
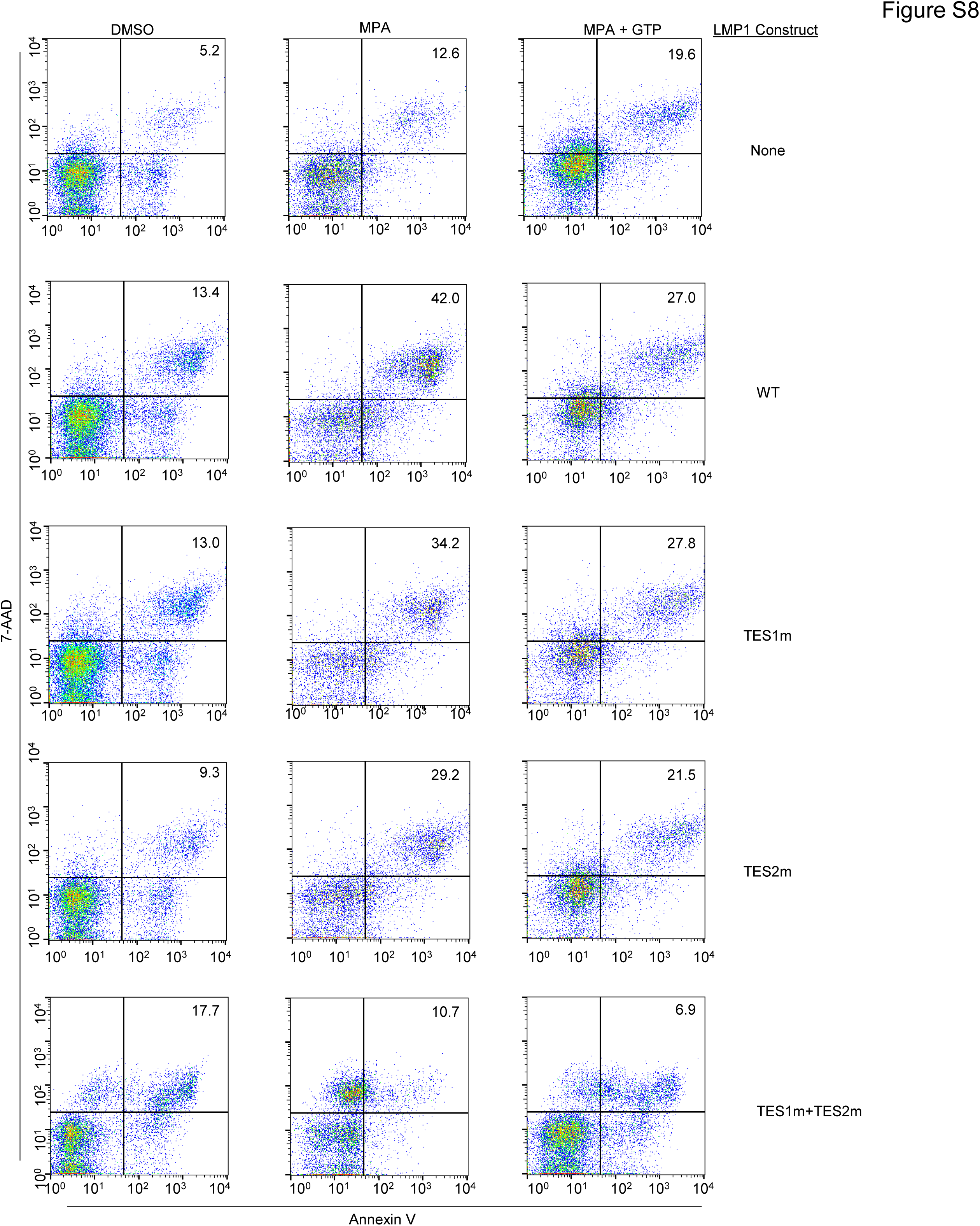
LMP1 TES1 and TES2 signaling sensitize EBV-negative BL41 Burkitt cells to MPA-driven apoptosis in a partially GTP dependent manner. Representative FACS plots from n=3 replicates of BL41 cells mock induced or induced for WT, TES1m, TES2m or TES1m+TES2m LMP1 expression for 24 hours and then treated with 1μM MPA ± 100μM GTP for 96 hours as indicated. Shown are FACS analysis of 7-AAD uptake versus Annexin V positivity.

**Figure S9.**
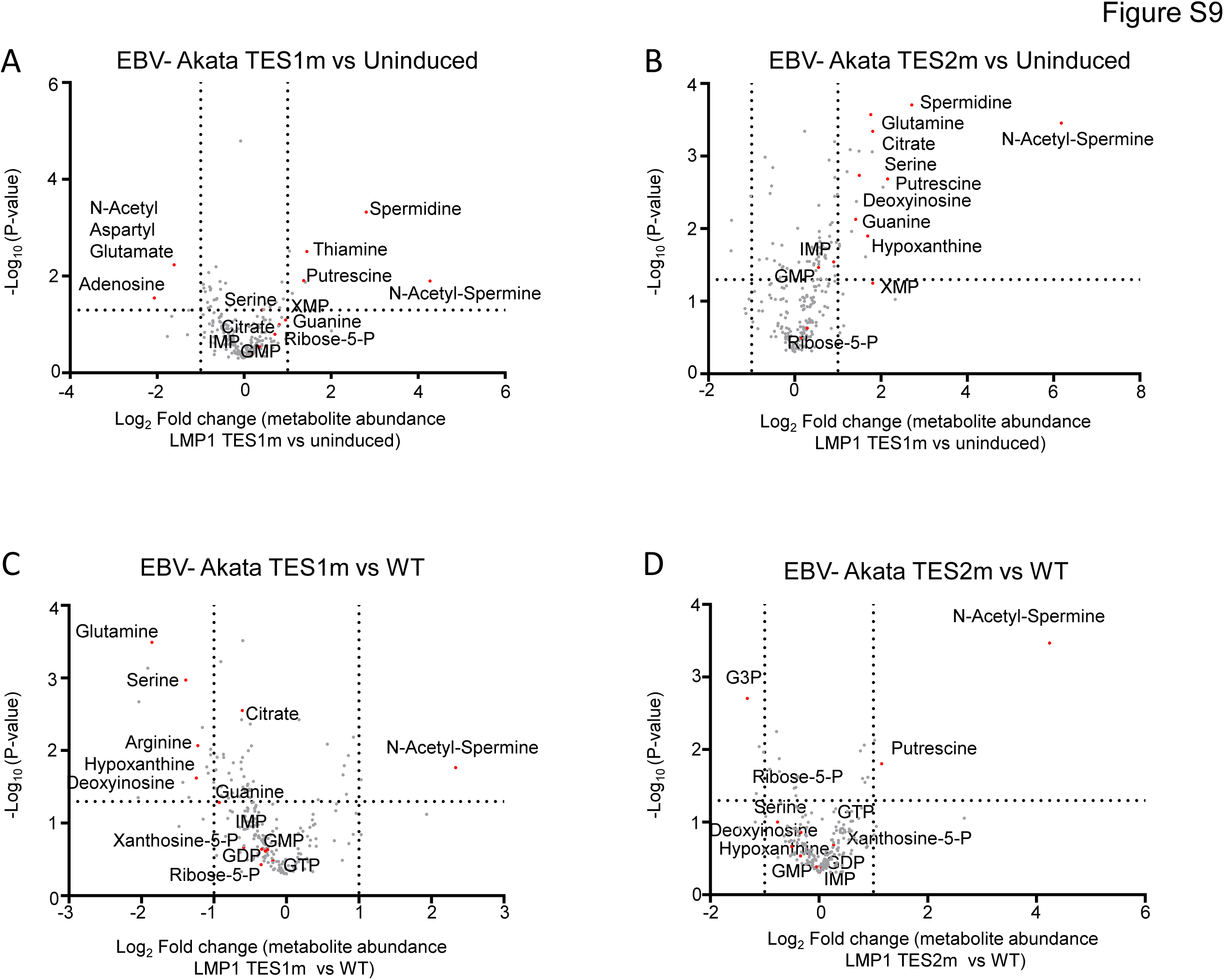
Effects of LMP1 TES1 versus TES2 signaling on Akata Burkitt metabolome remodeling. (A) Volcano plot of LC-MS metabolomic analysis of n=6 replicates of EBV-negative Akata cells mock induced or doxycycline induced for LMP1 TES1m expression for 24 hours. Metabolites with higher abundance in LMP1 TES1+ cells have positive fold change values, whereas those higher in mock induced cells have negative fold change values. Selected metabolites are highlighted by red circles and annotated. (B) Volcano plot of LC-MS metabolomic analysis of n=6 replicates of EBV-negative Akata cells mock induced or doxycycline induced for LMP1 TES2m expression for 24 hours, with selected metabolites highlighted as in (A). (C) Volcano plot of LC-MS metabolomic analysis of n=6 replicates of EBV-negative Akata cells doxycycline induced for TES1m vs WT LMP1 expression for 24 hours, with selected metabolites highlighted. Replicates for this cross-comparison were induced side by side, prepared for and analyzed by LC-MS together on the same day to minimize batch effects. (D) Volcano plot of LC-MS metabolomic analysis of n=6 replicates of EBV-negative Akata cells doxycycline induced for TES2m vs WT LMP1 expression for 24 hours, with selected metabolites highlighted. Replicates for this cross-comparison were induced side by side, prepared for and analyzed by LC-MS together on the same day to minimize batch effects.

**Figure S10.**
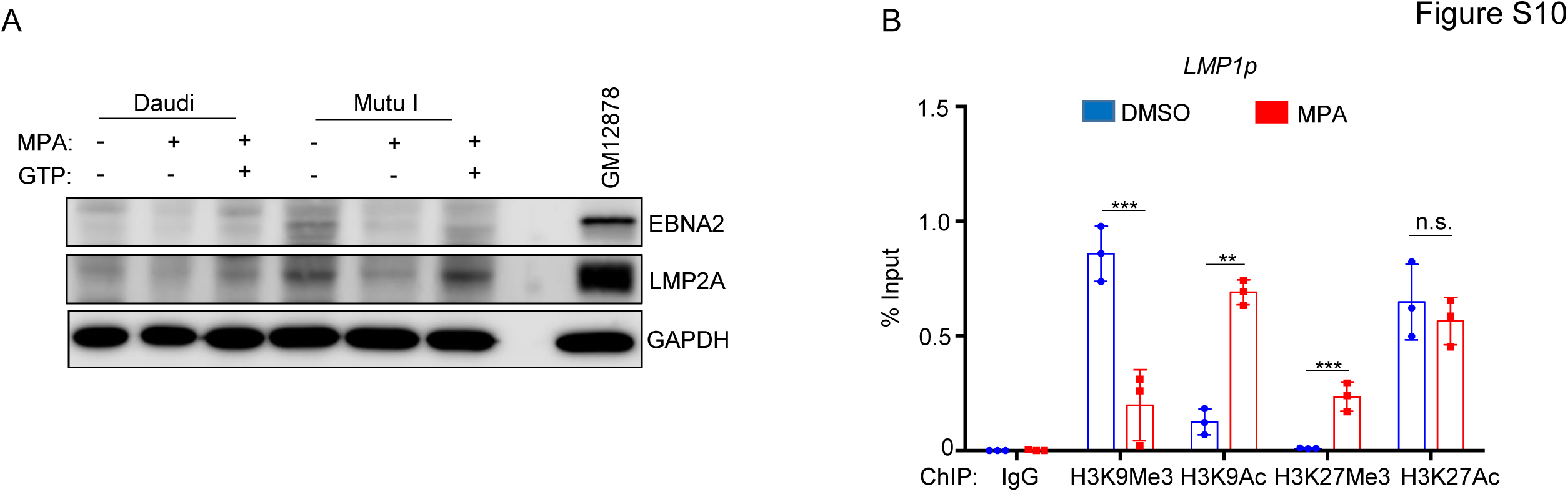
MPA does not de-repress EBNA2 or LMP2A latency III protein expression in Burkitt cells. (A) Immunoblot analysis of WCL from Daudi (left) versus MUTU I (right) treated with DMSO, 1µM MPA ± 100µM GTP for 24 hours. WCL for latency III GM12878 LCls was run in the rightmost lane as a positive control for EBNA2 and LMP2A expression. Daudi contain an EBV genomic deletion that knocks out the EBNA2 gene. (B) ChIP-qPCR analysis of MUTU I cells treated with DMSO or 1 μM MPA for 72 hours, using the indicated ChIP antibodies and qPCR primers specific for the LMP1p region.

